# Multiplexed mRNA assembly into ribonucleoprotein particles plays an operon-like role in the control of yeast cell physiology

**DOI:** 10.1101/2020.06.28.175851

**Authors:** Rohini R. Nair, Dimitry Zabezhinsky, Rita Gelin-Licht, Brian Haas, Michael C.A. Dyhr, Hannah S. Sperber, Chad Nusbaum, Jeffrey E. Gerst

## Abstract

Prokaryotes utilize polycistronic messages (operons) to co-translate proteins involved in the same biological process. Whether eukaryotes achieve similar regulation by selectively assembling monocistronic messages derived from different chromosomes is unclear. We employed transcript-specific RNA pulldowns and RNA-seq/RT-PCR to identify mRNAs that co-precipitate into ribonucleoprotein (RNP) particles in yeast. Consistent with the hypothesis of eukaryotic RNA operons, mRNAs encoding components of the mating pathway, heat shock proteins, and mitochondrial outer membrane proteins multiplex *in trans* to form discrete mRNP particles, termed transperons. Chromatin-capture experiments reveal that genes encoding multiplexed mRNAs physically interact, thus, RNA assembly may result from co-regulated gene expression. Transperon assembly and function depends upon H4 histones and their depletion leads to defects in RNA multiplexing, resulting in decreased pheromone responsiveness and mating, and increased heat shock sensitivity. We propose that intergenic associations and non-canonical H4 histone functions contribute to transperon formation in eukaryotic cells to regulate cell physiology.

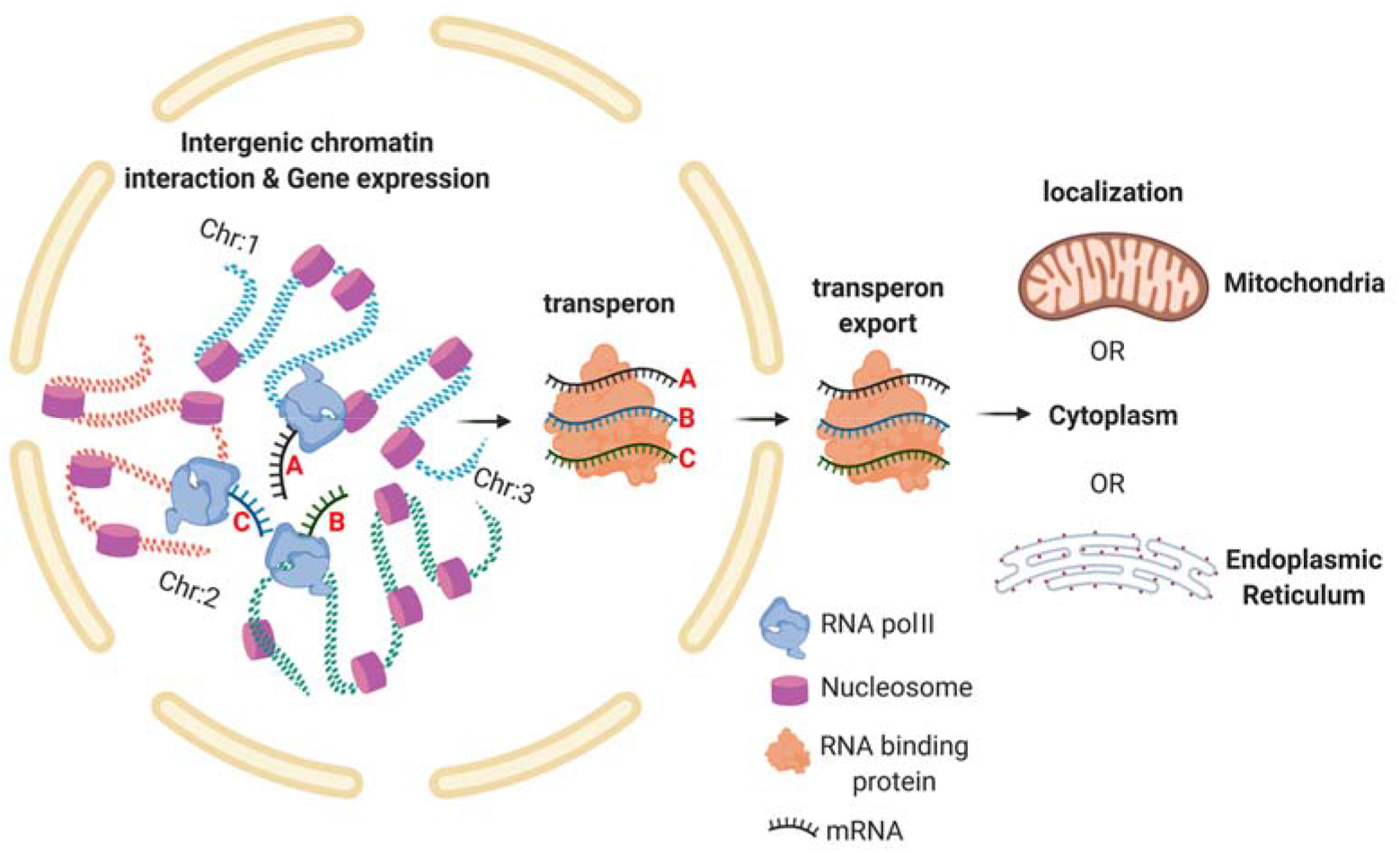

## Introduction

Prokaryotic organisms can rely on polycistronic transcription (operons), which allows for the expression of needed mRNAs from a single promoter and enables a rapid and robust response to corresponding stimuli (*e.g*. lactose operon) (Jacob and Monod, 1961). However, this mode of action has not been reported in eukaryotes, except for a few limited examples (*e.g*. in *C. elegans*) (Spieth et al., 1993), and eukaryotic operons have been observed to give rise to dicistronic messages (Blumenthal, 2004). However, it is intriguing to postulate whether eukaryotes have devised functional alternatives to operons. One possibility is that eukaryotic messenger ribonucleoprotein (mRNP) particles, which are composed of multiple RNAs and RNA-binding proteins (RBPs) (Mitchell and Parker, 2014), effectively confer combinatorial gene expression networks similar to prokaryotic operons (Keene and Tenenbaum, 2002). In this case, however, the mRNP particles contain individual mRNAs that undergo co-translational control and may encode proteins involved in the same biological process or molecular complex. The idea that mRNPs might essentially constitute RNA operons (Keene, 2007) or *transperons*, as termed here, implies several features. First, although mRNAs derived from different chromosomes can reside in the same mRNP, they should have common regulatory elements not only for transcriptional and translational control, but also for interacting at the post-transcriptional level with other mRNAs within the particle. The latter include motifs/structures to facilitate interactions with shared RBPs, as well as elements that might facilitate RNA-RNA interactions (*e.g*. base-pairing). Second, some mechanism must confer the recruitment and assembly *in trans (i.e*. multiplexing) of the individual mRNAs into single mRNP particles. Third, in order to be functionally relevant, transperons should contain mRNAs encoding proteins involved in the same functional context (*e.g*. organelle biogenesis, macromolecular complex, or biological process). Although the mechanism remains unknown, studies have shown that genomic DNA folds create sites of transcriptional hot spots (Lieberman-Aiden et al., 2009; Rao et al., 2014) that could facilitate multiplexing by co-localizing messages during transcription and coordinating the subsquent association of RBPs. Moreover, RNA-RNA interactome studies show that extensive interactions can occur directly between RNAs in living cells (Aw et al., 2016; Engreitz et al., 2014; Kudla et al., 2011; Nguyen et al., 2016; Sharma et al., 2016). Thus, the multiplexing of mRNAs into mRNPs to yield transperons should be directly testable.

Although individual yeast RBPs have been shown to bind to numerous (*e.g*. 10’s-1000’s of mRNAs) (Hogan et al., 2008), these RNA-binding studies are based primarily upon cross-linking and RBP pulldowns that are unable to define the minimal number of mRNA species in a single mRNP particle. To test the idea of RNA multiplexing and define the composition of such particles, we employed a single mRNA species pulldown procedure (RaPID) (Slobodin and Gerst, 2010; Slobodin and Gerst, 2011). This method employs MS2 aptamer-tagging of endogenously expressed messages in yeast (Haim et al., 2007) and their precipitation from cell lysates via the MS2 coat protein (MCP), to identify bound non-tagged transcripts using RNA-seq (RaPID-seq). We previously demonstrated that RaPID-seq identifies MS2-tagged target mRNAs (Haimovich et al., 2016) and now show that additional messages associate with these transcripts. Using *MAT*α yeast, we found that the pulldown of several target mRNAs (*e.g. SRO7, EXO70, OM45*) co-precipitated mRNAs that encode secreted proteins involved in cell mating (*e.g. STE3, SAG1, MF*α*1, MF*α*2*). Since a corresponding complex (*e.g. STE2, AGA1, AGA2, MFA1, MFA2*) could also be precipitated from *MAT*a cells, it demonstrated that mRNAs encoding secreted proteins involved in mating (*e.g*. pheromones, pheromone receptors, agglutinins, and proteases) multiplex into a functionally selective mRNP particle, the mating transperon. To identify factors involved in mating transperon assembly, pulldowns of these mRNAs followed by mass spectrometry (RaPID-MS) revealed that the yeast histone H4 paralogs, Hhf1 and Hhf2 (Dollard et al., 1994), interact with the mRNAs to regulate particle assembly. Furthermore, both pheromone responsiveness (*e.g*. shmooing) and mating were inhibited by either histone H4 depletion or block in acetylation. No other histones when mutated had this effect. Conserved *cis* elements in the mating mRNAs were also identified and mutations therein had deleterious effects upon mRNA multiplexing, pheromone responsiveness and mating, similar to those seen upon histone H4 mutation.

To determine the mechanism by which mRNA multiplexing might occur, we performed chromatin conformation capture (Lieberman-Aiden et al., 2009), as well as genomic locus tagging experiments, to look for evidence of direct allelic interactions. A specific interaction between the *STE2* and *AGA2* genes was identified in *MAT*a cells, which suggests that interallelic coupling may give rise to RNA multiplexing. To confirm this hypothesis, we examined whether mRNAs encoding heat shock proteins (HSPs), whose genes are already known to undergo intergenic interactions during heat shock (Chowdhary et al., 2019), undergo multiplexing. Importantly, HSP mRNAs also multiplex *in trans* during heat shock to form RNP particles and a specific requirement for histone H4 was shown to facilitate survival afterwards. Parallel studies revealed the existence of additional RNA multiplexes for mitochondrial outer membrane proteins and MAP kinase pathway proteins in yeast. Overall, these results suggest that histone H4-mediated chromatin interactions in eukaryotic cells lead to the formation of functionally selective mRNP particles, as a potential means to co-regulate gene expression in place of polycistronicity.

## Results

### mRNAs encoding yeast mating pathway components co-precipitate in mRNPs

We developed the RaPID procedure (Slobodin and Gerst, 2010; Slobodin and Gerst, 2011) to identify RBPs and mRNAs that bind to specific RNAs of interest. Unlike CLIP or PAR-CLIP, which collect information on all RBP-RNA interactions in the cell and is biased toward highly expressed mRNAs with long poly-A tails (Hafner et al., 2010), RaPID allows for a biochemical view of the protein and RNA constituents of mRNPs at the single transcript level. RaPID employs the precipitation of MS2 aptamer-tagged messages using the MS2 coat protein (MCP) fused to streptavidin-binding peptide, followed by elution from immobilized streptavidin with free biotin, and then mass spectometry (Slobodin and Gerst, 2010; Slobodin and Gerst, 2011) or RNA-seq (Haimovich et al., 2016) to identify bound proteins and RNAs, respectively. In order to address issues regarding whether the MS2 system might affect the stability of MS2 aptamer-tagged messages, we performed RaPID and RNA-seq (RaPID-Seq) on eleven endogenously expressed MS2 aptamer-tagged messages in *MAT*α yeast (Haimovich et al., 2016). These included representative transcripts having a wide range of intracellular patterns of localization [*e.g*. mitochondria: *OXA1* and *OM45*; cortical endoplasmic reticulum (cER)/asymmetrically-localized mRNAs: *ABP1, SRO7, EXO70, ASH1, MYO2*, and *MYO4*; peripheral nuclear endoplasmic reticulum (nER): *ATG8*; and peroxisomes: *PEX14*] (Gadir et al., 2011; Haim et al., 2007; Zipor et al., 2009). In addition to identifying the target mRNA in each pulldown, we identified non-tagged RNAs that co-precipitated with the targets (Figures 1A and S1, and see Table S1 for all RNA-seq data). These included retrotransposable elements (*e.g*. Ty elements), tRNAs, ribosomal RNAs, telomere and small nuclear RNAs (Figure S1), and coding RNAs (Figures 1, S1, and Table S1). Notably, five mRNAs encoding secreted and membrane proteins (mSMPs) of the mating process were found associated with a subset of target RNAs. These target transcripts consisted of two cER-localized mRNAs that encode polarity factors (mPOLs; *SRO7* and *EXO70*) (Aronov et al., 2007) and *OM45*, an mRNA that encodes a mitochondrial outer membrane protein (MOMP), that we have shown to localize to both mitochondria and ER (Gadir et al., 2011). Co-precipitated non-tagged mRNAs included *STE3*, which encodes the a-pheromone receptor, *MF*α*1* and *MF*α*2*, which both encode α-mating factor, *SAG1*, which encodes α-agglutinin, and *AFB1*, that encodes the a-pheromone blocker (Figure 1B). This result is consistent with the idea that specific RNAs could multiplex to form a discrete mRNP particle.

**Figure 1.**
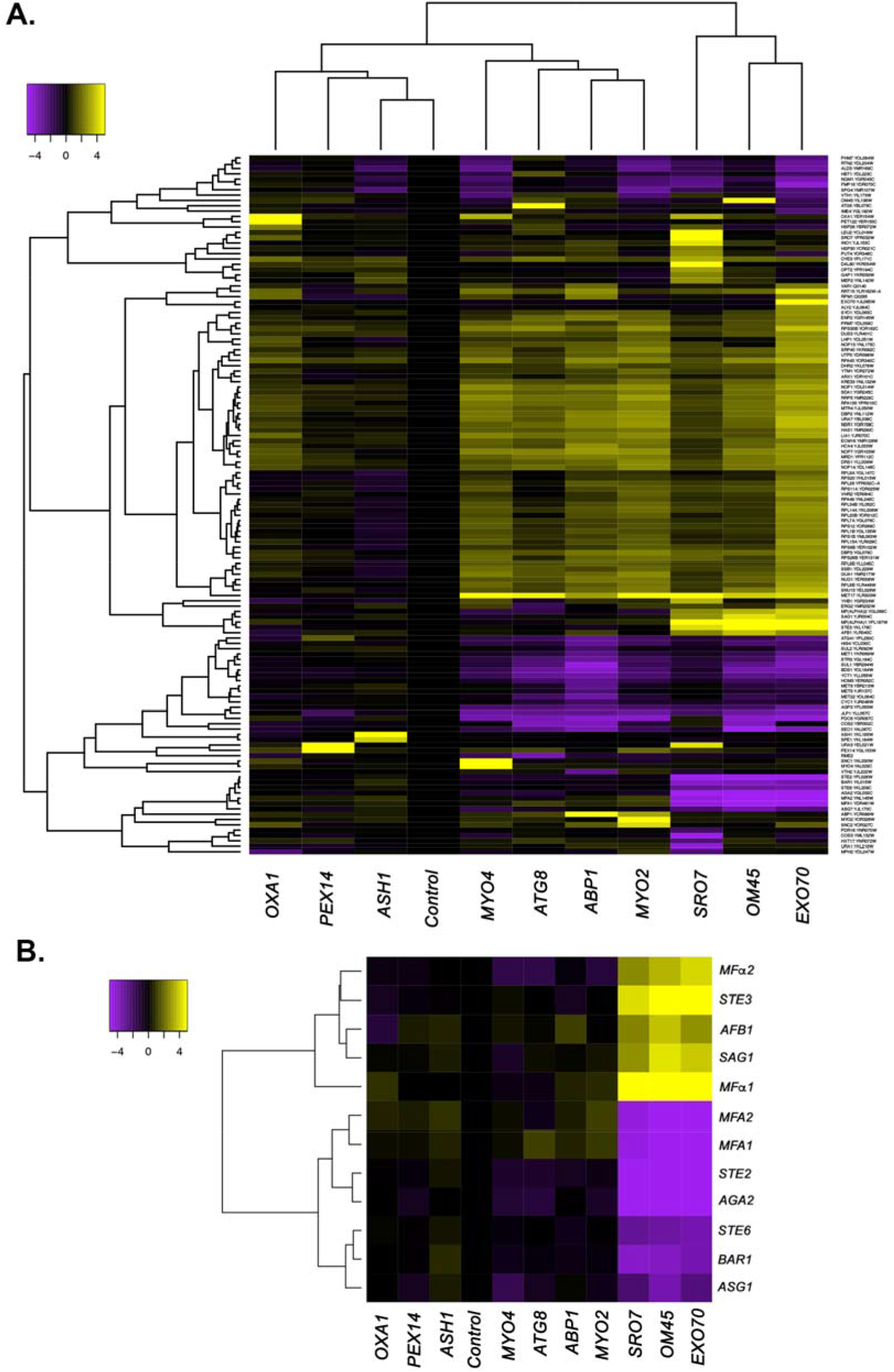
mRNAs encoding secreted and membrane proteins involved in yeast mating multiplex into a single ribonucleoprotein particle. (A) RaPID RNA pulldown and RNA-seq reveals transcript multiplexing in yeast. Different MS2 aptamer-tagged mRNAs (listed on x axis) expressed from their genomic loci were precipitated from *MAT*α yeast (BY4742) by MS2-CP-GFP-SBP after formaldehyde crosslinking *in vivo* and cell lysis procedures (RaPID; see Materials and Methods). RNA-seq was performed and the reads were plotted to yield a heat-map for the relative enrichment of the tagged as well as non-tagged mRNAs (see color key for approximate values). All non-coding RNAs (*e.g*. Ty elements, tRNAs, snoRNAs, ribosomal RNAs) were removed, but are included in the heatmap of Supplementary Figure S1. A list of the log2 transformed and control-subtracted expression values, as plotted here, is given in Table S1. (B) mRNAs encoding proteins involved in mating in *MAT*α cells are enriched in the pulldowns of *SRO7, EXO70*, and *OM45* mRNAs. A selective heat map comprising the mRNAs encoding secreted and membrane proteins involved in yeast cell mating is shown. Positive enrichment of cell-type specific *MAT*α mRNAs (*STE3, MF*α*1, MF*α*2, SAG1, AFB1*) is shown in purple, while negative enrichment of the non-expressed *MAT*a mRNAs (*MFA1, MFA2, STE2, AGA2, BAR1, STE6, ASG7*) is shown in yellow.

Importantly, the corresponding *MAT*a mating-type RNAs (*e.g. STE2, MFA1, MFA2, AGA2*, and *BAR1*), as well as other *MAT*a RNAs involved in mating (*e.g. STE6, ASG7*), were negatively enriched (depleted) in these pulldowns (Figure 1A and B). While this result was predicted, due to their lack of expression in *MAT*α cells, nonetheless it validates the specific pulldown of the *MAT*α mating mRNAs.

We verified the RaPID-Seq results using RaPID followed by RT-PCR (RaPID-PCR) and found that the pulldown of *SAG1* mRNA co-precipitated its cohort mRNAs of the mating pathway (*e.g. STE3, MF*α*1, MF*α*2*), but not unrelated mRNAs (*e.g. UBC6, SEC3, WSC2*) (Figure 2A). We noted that the original target mRNAs (*EXO70, SRO7, OM45*) were present in this pulldown, though not as highly represented in the RT-PCR data as those mRNAs directly related to mating. We note that *IST2* mRNA was identified in the pulldown from *MAT*α cells (Figure 2A), but not from *MAT*a cells (see below; Figure 2B).

**Figure 2.**
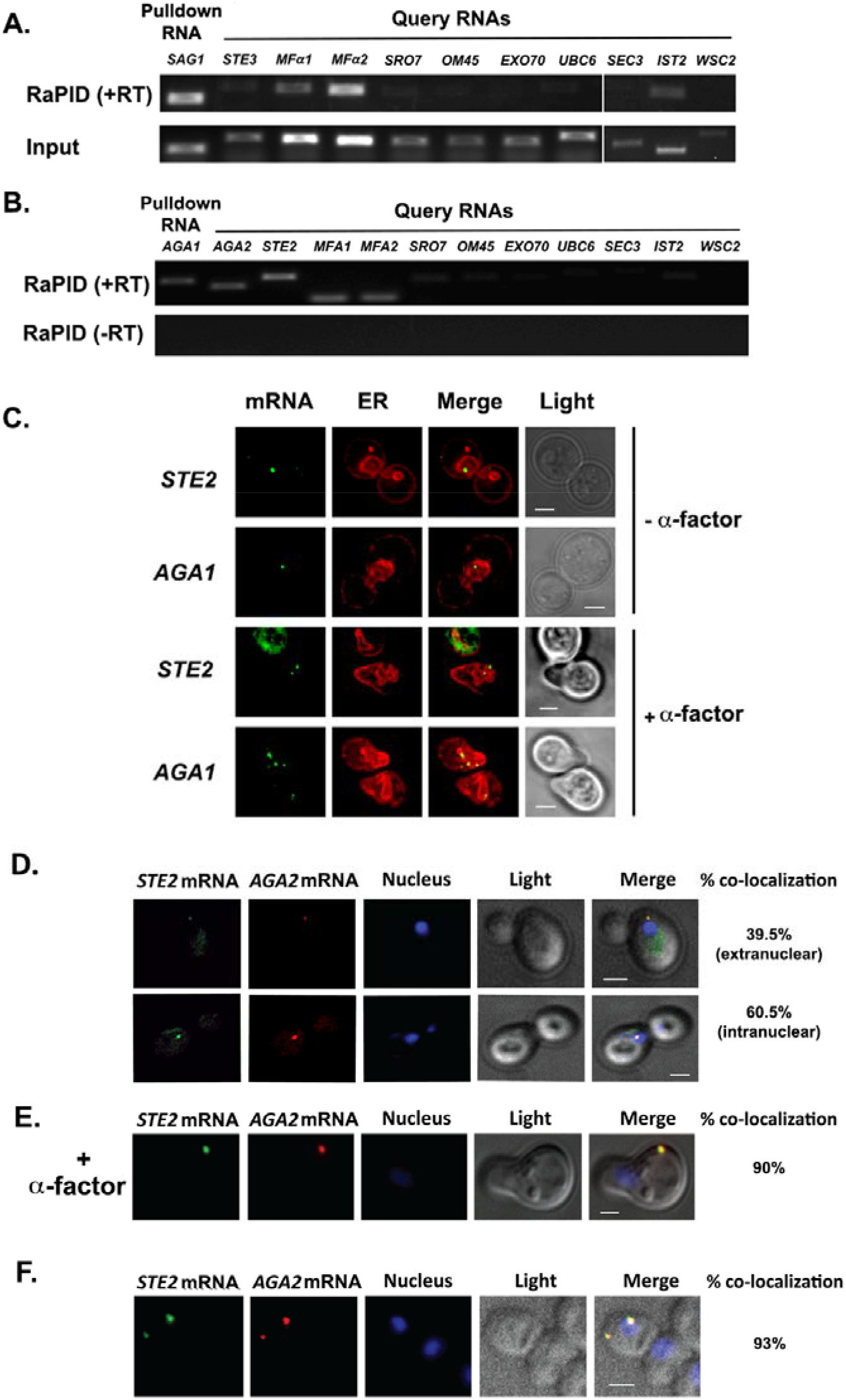
mRNAs encoding secreted and membrane proteins involved in mating multiplex into a ribonucleoprotein particle in both *MAT*α and *MAT*a cells and localize to nER. (A). A mating mRNP is also formed in *MAT*a cells. *MAT*a yeast (BY4741) expressing MS2 aptamer-tagged *SAG1* from its genomic locus were grown to mid-log phase (O.D._600_=0.5) and subjected to RaPID followed by RT-PCR (RaPID-PCR; see Materials and Methods). RNA derived from the total cell extract (Input) or biotin-eluated fraction (RaPID) was analyzed by RT-PCR using primer pairs corresponding to mRNAs expected to possibly multiplex (based upon Figure 1). (B) The mating mRNP in *MAT*α cells can be analyzed by RaPID-PCR. *MAT*α yeast expressing MS2 aptamer-tagged *STE2* from its genomic locus were grown and processed as in *A*. RNA derived from the biotin-eluated fraction (RaPID) was analyzed by PCR either with (+RT) or without (-RT) reverse transcription using primer pairs corresponding to mRNAs expected to multiplex (based upon Figure 1). (C) Mating mRNAs localize to nER in pheromone-treated or non-treated cells. Wild-type cells expressing MS2 aptamer-tagged *STE2* or *AGA1* mRNAs (as listed) were transformed with single-copy plasmids expressing MS2-CP-GFP(x3) and Scs2-mCherry (an ER marker). Cells were grown to mid-log phase and either α-factor (5μM; + α-factor) or DMSO (-α-factor) was added for 90 minutes, followed by fixation with 4% paraformaldehyde in medium containing 3.5% sucrose, and prior to imaging by confocal microscopy. *mRNA* - MS2-CP-GFP(x3) labeling; *ER* – (Scs2-mCherry labeling); *merge* – merger of mRNA and ER windows; *light* – transmitted light. Size bar = 2μm. (D) Mating mRNAs *STE2* and *AGA2* co-localize both in nucleus and outside of the nucleus. Yeast expressing MS2 aptamer-tagged *STE2* mRNA and PP7 aptamer tagged *AGA2* mRNA from its genomic locus were transformed with MS2-CP-GFP(x3) and PP7-PS-tomato, while the nucleus was stained with DAPI. *STE2 mRNA* - MS2-CP-GFP(x3) labeling; *AGA2 mRNA* – PP7-PS-tomato labeling; Nucleus - DAPI labeling; *merge* – merger of mRNA and nucleus windows; *light* – transmitted light. Size bar = 2μm. (E) Mating mRNAs *STE2* and *AGA2* co-localize upon pheromone treatment. Cells used in (*D*) were treated with α-factor (10μM) and visualized after 3-4hr. Nucleus - DAPI labeling; *merge* – merger of mRNA fluorescence and DAPI windows; *light* – transmitted light. Size bar = 2μm. (F) smFISH validation of *STE2* and *AGA2* mRNA co-localization. Untransformed cells from (*D*) were processed for smFISH labeling using non-overlapping FISH probes complementary to the MS2 and PP7 aptamers, prior to labeling with DAPI. mRNA granule scoring was performed using the FISH-quant algorithm and co-localization of the granules was performed manually. Size bar= 2μm.

By employing RaPID-PCR, we next examined whether a corresponding set of mating pathway mRNAs [*i.e. STE2* (α-factor receptor), *AGA1* and *AGA2* (agglutinins), and *MFA1* and *MFA2* (a-factor)] could co-precipitate in *MAT*a cells. Indeed, the precipitation of *AGA1* mRNA using RaPID led to co-precipitation of the cohort mating pathway mRNAs (*e.g. STE2, AGA2, MFA1, MFA2*; Figure 2B), along with some *SRO7* and *OM45*. Thus, we demonstrated the existence of a mRNP particle for the mating pathway of both haplotypes.

### mRNAs encoding mating pathway components co-localize to nER in both vegetative-growing and pheromone-treated cells

We examined whether mRNAs of the mating mRNP particle localize to the nER, like other mSMPs (Kraut-Cohen et al., 2013) or, perhaps to the bud/shmoo tip like asymmetrically localized mRNAs that encode polarity and secretion factors (Aronov et al., 2007; Gelin-Licht et al., 2012). A previous study suggested that aptamer-tagged mating pathway mRNAs, like *MFA2*, localize to P-bodies located in (or near) the shmoo tip and that these granules are functional sites for transmitting the mating signal (Aronov et al., 2015). However, a more recent work by us (Haimovich et al., 2016) showed that endogenously expressed MS2 aptamer-tagged *MFA2* mRNA does not localize to P-bodies and that P-bodies only form under conditions of mRNA over-expression, as in Aronov *et al*. (2015). We examined the localization of endogenously expressed MS2-tagged *STE2* and *AGA1* mRNAs, along an ER marker, mCherry-Scs2, in both non-treated and α-factor (1μM)-treated cells. We found that both mRNAs co-localize with nER under both growth conditions and are not present either in the bud or shmoo tips (Figure 2C). In addition, we examined the localization of endogenously expressed MS2 aptamer-tagged *MFA1* and *MFA2* mRNA in wild-type cells or cells lacking *SHE2*, an ER-localized RBP involved in the asymmetric localization of mRNAs to the bud (but not shmoo) tip (Genz et al., 2013). Both *MFA1* and *MFA2* mRNAs localized to the cell bodies in either pheromone-treated and non-treated cells, and no difference in localization was observed upon the deletion of *SHE2* (Figure S2A). Thus, like other mSMPs, RNAs of the mating particle are not trafficked to the polarized extensions of yeast cells (Kraut-Cohen et al., 2013), nor localize to P-bodies as previously suggested (Aronov et al., 2015).

As RNA co-localization is predicted by RNA multiplexing, we tagged endogenous *AGA2* with the PP7 aptamer (Larson et al., 2011) in the cells expressing MS2-tagged *STE2* and examined for co-localization upon expression of their respective fluorescent protein-tagged aptamer-binding proteins (*e.g*. PP7-PS-tomato and MS2-CP-GFP(x3)). Overall, we found that 65.5+7.7% (average + standard deviation; n = 3 experiments) of PP7-tagged *AGA2* mRNA co-localized with MS2-tagged *STE2* mRNA (Figure 2D), of which 39.5+3.5% colocalized in the cytoplasm and 60.0+3.5% co-localized in the nucleus. Importantly, the stimulation of cells with α-factor (10μM) increased the overall co-localization of *AGA2* mRNA with *STE2* mRNA to 90.0+7.1% (Figure 2E). We validated the live imaging results using single-molecule fluorescence *in situ* hybridization (smFISH). Specific probes against the MS2 and PP7 aptamers were used to score the localization of *STE2* and *AGA2* mRNAs, respectively. The number of FISH spots per cell was quite variable for either mRNA, *e.g*. the average number of spots (RNA molecules) for *STE2* mRNA was 6.4+4.3, while that of *AGA2* mRNA was 1.1+1.0. Nevertheless, 92.9+1.2% of PP7-tagged *AGA2* mRNA spots co-localized with MS2-tagged *STE2* mRNA spots, and of these, 38.2+2.5% co-localized in the cytoplasm, whereas 61.8+2.5% co-localized in the nucleus (Figure 2F). Thus, mating mRNAs shown to multiplex using biochemical means also can be shown to co-localize before and after nuclear export.

### A histone H4 paralog binds to mating mRNP RNAs

To determine how the mating mRNP particle assembles, we performed RaPID-MS using the *MFA1, MFA2*, and *STE2* mRNAs either bearing or lacking their 3’UTRs, along with an unrelated control RNA, *ASH1*, as target RNAs. We identified a band present at ~150kDa that was present in all lanes upon silver staining, except for the lanes corresponding to *STE2* mRNA lacking its 3’UTR and *ASH1* mRNA (Figure S2B). Sequencing revealed that this band contained a core histone H4 paralog, Hhf1. Although the predicted molecular mass of Hhf1 is ~11kDa, the likelihood exists that it was cross-linked with another protein. Indeed, other cross-linked products were found in this band although none were shown to be connected to mRNAs of the mating mRNP.

To verify whether Hhf1 alone can precipitate the mating mRNP particle, we performed immunoprecipitation using HA-tagged Hhf1 and examined the precipitates for mating mRNAs by RT-PCR. We found that Hhf1 could precipitate either native or MS2 aptamer-tagged *STE2*, along with native *MFA1* or *MFA2* mRNA (Figure 3A). In contrast, we could not detect the presence of *UBC6, ASH1, EXO70*, or *SRO7* mRNA in these pulldowns (Figure 3B). Thus, we could verify specific interactions between Hhf1 and RNAs of the mating particle. In addition, we could show that Hhf1 migrates at around 12kDa in SDS-PAGE gels (Figure 3C), as predicted.

**Figure 3.**
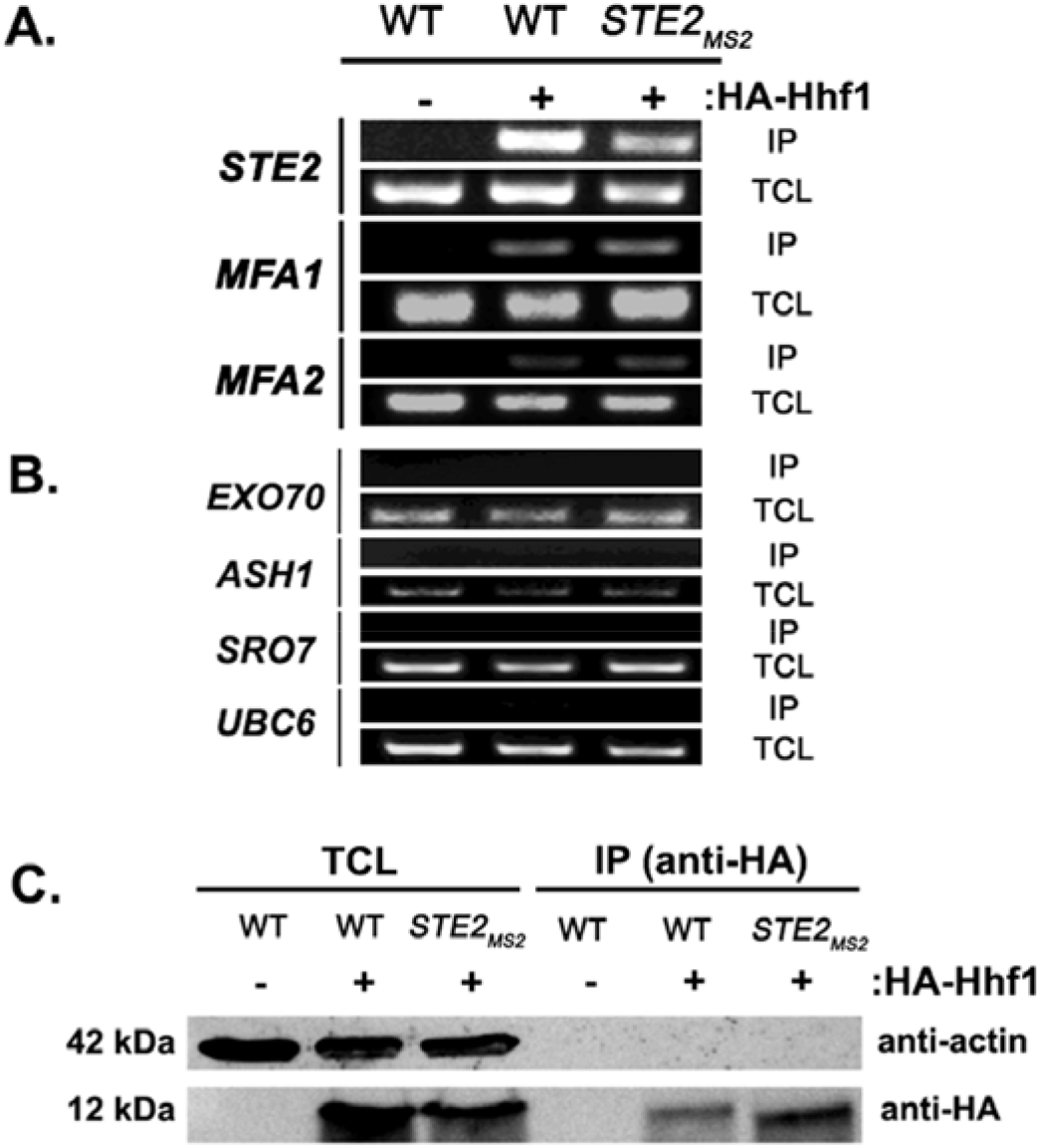
HA-Hhf1 precipitates the mating mRNP. (A & B) HA-Hhf1 precipitates the mating mRNAs. *MAT*a wild-type (WT) cells and cells expressing MS2 aptamer-tagged *STE2* mRNA from its genomic locus were transformed with a 2μ plasmid expressing *HA-HHF1* (+ HA-Hhf1). WT cells were also transformed with an empty vector (-HA-Hhf1), as control. Cells were grown to mid-log phase, lysed to yield the total cell lysate (TCL) and subjected to immunoprecipitation (IP) with anti-HA antibodies. Following IP, the RNA was extracted, DNase-treated, and reverse-transcribed. Transcripts were amplified using specific primers to the genes listed. A representative ethidium-stained 1% agarose gel of electrophoresed samples is shown; n=3 experiments. (C) HA-Hhf1 migrates as a 12kDa protein in SDS-PAGE gels. Aliquots (5% of total) of the TCL and IP fractions were resolved on 15% SDS-PAGE gels and transferred to nitrocellulose membranes for detection with anti-HA and actin antibodies to verify the presence of HA-tagged Hhf1 in both fractions.

### Histone H4 depletion affects mating

Although histones are principally known for their DNA-binding functions in nucleosome assembly (Wu and Grunstein, 2000), we determined whether Hhf1 or its paralog, Hhf2, play a role in mating RNP assembly and mating. First, we examined whether deletions in H4 or other histone genes affect mating. We performed quantitative mating assays by crossing wild-type (WT) yeast against single gene mutations in the H2A, H2B, H3, and H4 paralog pairs. We found that only deletions in H4 (*hhf1Δ or hhf2Δ*) led to a significant decrease (~50-60%) in mating (Figure 4A). Since histone paralog pairs are essential, we created a conditional allele of *HHF1* at its genomic locus by adding an auxin-induced degron sequence at the 5’ end (*HA-AID-HHF1*) along with an HA epitope, in *hhf2Δ* cells. This allele had no effect upon growth in the absence of auxin (3-indole acetic acid; 3-IAA), but led to lethality upon long-term treatment in the presence of 3-IAA (4mM) (Figure S3A). Growth of the cells in presence of 3-IAA resulted in a full depletion of Hhf1 within 4hrs (Figure S3B) and cells remained viable after this short-term treatment (Figure S3C). When examined for mating after pre-treatment with 3-IAA (4mM; 4hrs), we observed a >80% inhibition in mating (Figure 4B). Thus, the depletion of histone H4 strongly affects yeast cell mating, even under conditions where viability is maintained.

**Figure 4.**
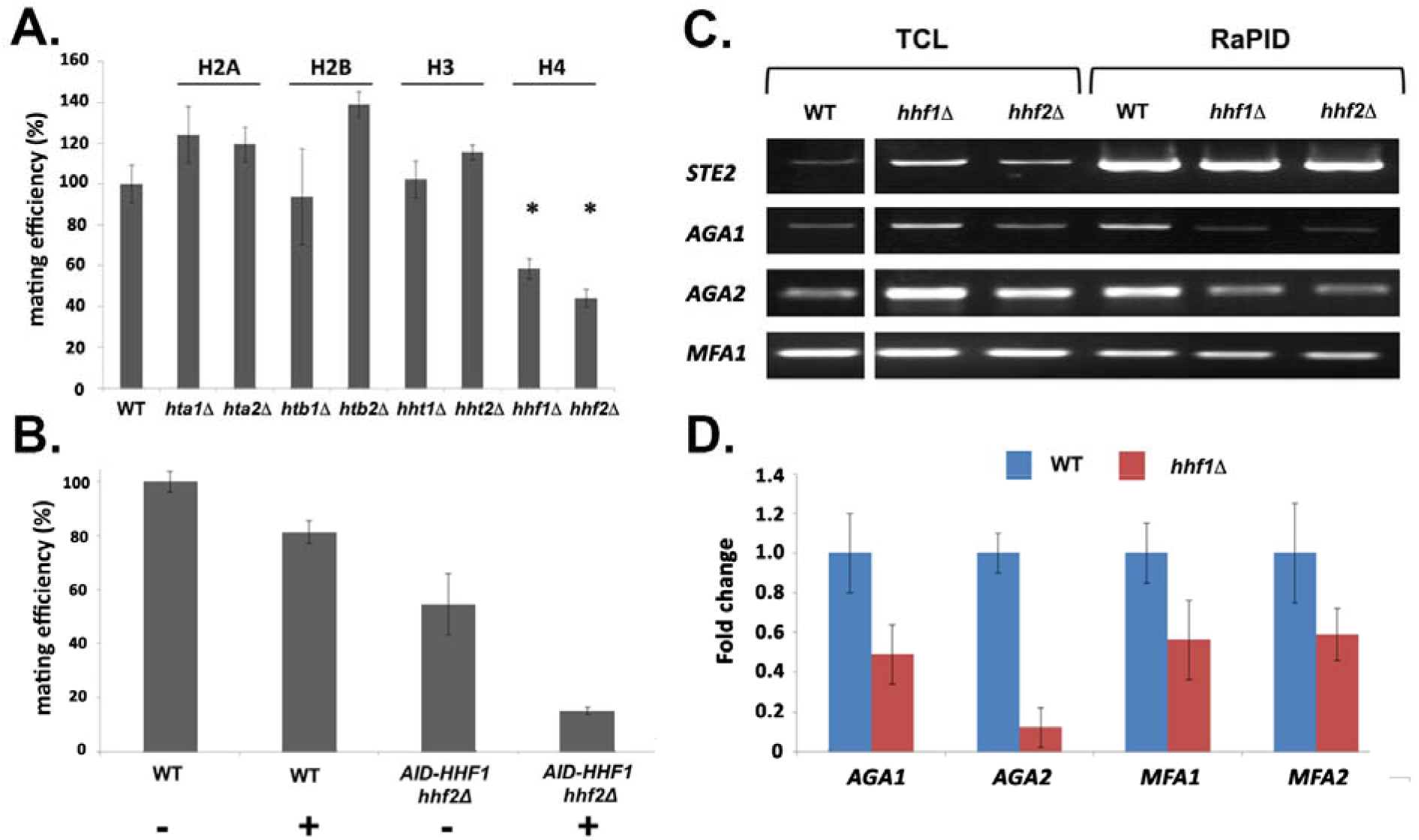
Histone H4 function is required for mating mRNP assembly and mating. (A) Histone H4 alone is required for mating. *MAT*a yeast (BY4741) bearing mutations in H2A (*hta1Δ, hta2Δ*), H2B (*htb1Δ, htb2Δ*), H3 (*hht1Δ, hht2Δ*), and H4 (*hhf1Δ, hhf2Δ*) were grown to mid-log phase, crossed against *MAT*α wild-type cells, and mating was scored using the quantitative mating assay. n = 3 experiments; * indicates p value <0.05 (B) Combined *hhf2Δ* and *AID-HHF1* alleles result in a near-complete block in mating upon auxin induction. *MAT*a wild-type (WT) and *hhf2Δ AID-HHF1* cells were grown to mid-log phase, treated either with or without auxin (3-IAA; 4mM) for 4hrs, and examined for mating using the quantitative mating assay. n = 3 experiments; *p* value <0.05. (C) Deletions in H4 histones result in defects in mating mRNP assembly – PCR analysis. *MAT*a WT, *hhf1Δ*, and *hhf2Δ* cells expressing MS2 aptamer-tagged *STE2* from the genome were grown to mid-log phase, processed for RaPID-PCR, and the extracted RNA analyzed using primers against the listed mating mRNAs. A representative ethidium-stained 1% agarose gel of electrophoresed samples is shown; n=3 experiments. (D) Deletions in a H4 histone result in defects in mating mRNP assembly – qPCR analysis. Same as in *C*, except that qRT-PCR was performed instead of PCR on WT and *hhf1Δ* cells expressing MS2 aptamer-tagged *STE2* from the genome. Three experiments were performed and gave similar results; *p* value <0.05.

To check whether the removal of H4 results in changes in the mating RNA levels, we examined *STE2* and *AGA1* transcript levels using qRT-PCR in WT and *AID-HHF1 hhf2Δ* cells either with or without added 3-IAA and/or α-factor. In the cases of *STE2* and *AGA1*, we observed similar increases in mRNA transcript levels in either cell type upon α-factor addition (Figure S3D) with or without added 3-IAA. Thus, auxin treatment does not abolish the increase in mating mRNA levels observed upon pheromone addition. In fact, we noted that auxin addition alone had a small stimulatory effect upon RNA levels (Figure S3D).

### Deletion of histone H4 alleles affects mating transperon assembly

As the depletion of H4 greatly lessens the mating propensity of yeast, we examined whether this is a consequence of altered mRNP particle assembly. We first performed RaPID-PCR on *STE2* mRNA derived from *hhf1Δ and hhf2Δ* cells, and measured the co-precipitation of other mating pathway mRNAs. We found decreased levels of *AGA1, AGA2*, and to a lesser degree *MFA1* mRNA in the *hhfΔ* deletion strains (Figure 4C). This result was verified using RaPID-qPCR where the levels of bound *AGA1, MFA1*, and *MFA2* mRNAs declined by ~50%, while that of *AGA2* declined even more (Figure 4D). The results indicate that the reduction in H4 amounts results in defects in mating mRNP formation, which may account for the loss in mating efficiency (Figure 4B).

### Histone acetyl transferases and histone H4 acetylation are required for mating

The N-terminus of histones is extensively modified by methylation, acetylation, and phosphorylation (Allis and Jenuwein, 2016). We examined the mating efficiency of yeast upon the deletion of three histone acetyl transferase genes (HATs; *e.g. HAT1, HAT2*, and *SAS2*) (Kurdistani and Grunstein, 2003) that were found to be physically and genetically linked with the *HHF1* gene and its product, according to the Saccharomyces Genome Database (SGD). We found that the deletion of *HAT1* or *SAS2* had the same reduced efficiency as observed with *hhf1Δ* cells (Figure 5A). This implies that the acetylated state of H4 is necessary for function.

**Figure 5.**
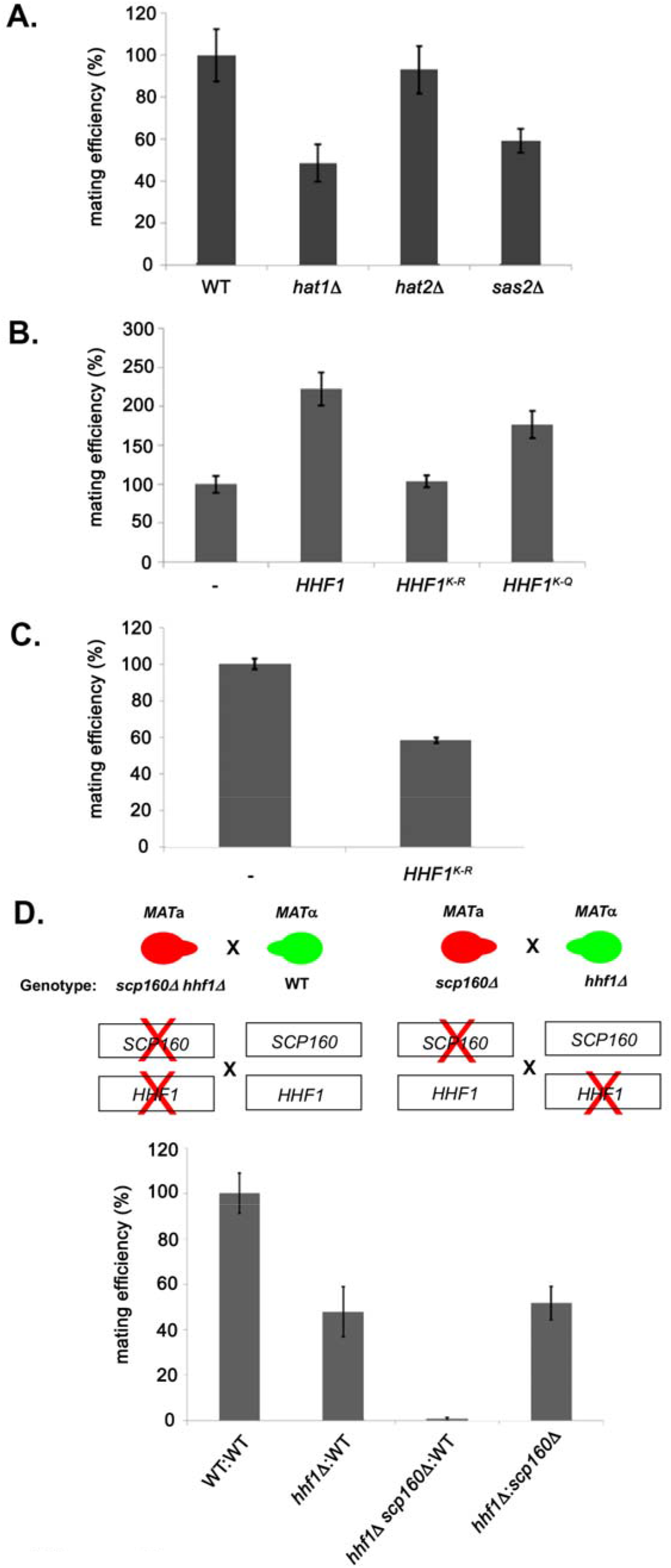
Histone H4 acetylation is required for mating. (A) Histone acetyltransferases are required for mating. Wild-type (WT; BY4741) and yeast lacking known acetyltransferase genes (*hat1Δ, hat2Δ*, and *sas2Δ*) were grown to mid-log phase (O.D._600_=0.5) and examined for mating with WT *MAT*α cells (BY4742). Three quantitative mating experiments were performed and gave similar results; *p* <0.05. (B) Mutations that either mimic or block N-terminal histone H4 acetylation increase or decrease mating efficiency, respectfully. *MAT*a wild-type (BY4741) cells alone (-) or over-expressing *HHF1* (*HHF1*), along with cells overexpressing *HHF1* bearing K-to-R (*HHF1^K-R^*) or K-to-Q (*HHF1^K-Q^*) mutations from a 2um plasmid plasmid were grown to mid-log phase and examined for mating with *MAT*α cells. Three experiments were performed; *p* <0.05. (C) Expression of the K-to-R mutation in *HHF1* in the genome inhibits mating comparative to the deletion of *HHF1. MAT*a wild-type (BY4741) cells alone (-) or expressing *HHF1* bearing the K-to-R (*HHF1^K-R^*) mutation from the genome were grown to mid-log phase and examined for mating with *MAT*α cells. Three experiments were performed; *p* <0.05. (D) *HHF1* and *SCP160* are epistatic and *hhf1Δ × Scp160Δ* crosses block mating. Crosses between wild-type (WT), *hhf1Δ*, or *hhf1Δ scp160 MAT*a cells and *MAT*α WT cells, along with an *hhf1Δ* and *scp160Δ* cross were performed. Illustrated are the *hhf1Δ scp160* × WT cross (upper left side) and the *hhf1Δ × scp160Δ* cross (upper right side). Three experiments were performed; *p* <0.05.

To verify this, we mutated all N-terminal Hhf1 lysine residues to arginines (*e.g*. K6R, K9R, K13R, K17R, K32R; K-to-R mutant) to abolish acetylation by HATs, or glutamine (*e.g*. K6Q, K9Q, K13Q, K17Q, K32Q; K-to-Q mutant) to mimic the acetylated state. We mated *MAT*a cells expressing these mutants with *MAT*α cells and assessed mating efficiency. Overexpression of WT *HHF1* caused a >2-fold increase in mating efficiency (Figure 5B), while the K-to-R mutant abolished it. Moreover, expression of the K-to-R mutant from the genome led to a 50% decrease in mating (Figure 5C). Correspondingly, overexpression of the K-to-Q mutant of Hhf1 yielded the same 2-fold increase in mating efficiency as observed upon *HHF1* overexpression (Figure 5B). Thus, histone acetylation is critical for H4 participation in the mating process.

### Histone H4 and Scp160 work cooperatively to confer mating

Previous work from the lab identified Scp160 as an ER-localized RBP involved in the delivery of certain mPOLs (*e.g. SRO7*) and mRNAs that confer the internal MAP kinase (MAPK) mating signal (*e.g. FUS3, KAR3, STE7*) to the shmoo tip upon pheromone treatment (Gelin-Licht et al., 2012). Deletion of *SCP160* was shown to prevent the polarized trafficking of these mRNAs on cER, resulting in a loss in chemotropism, a 60-70% decrease in mating in heterozygous crosses, and a >98% decrease in homozygous crosses. As there are two distinct ER paths (nER vs. cER) involved in localization of the two sets of mRNAs involved in mating (those encoding secreted components involved in the extracellular signaling aspect of mating, as mediated by Hhf1, and those encoding components of the MAPK cascade involved in the intracellular signaling aspect of mating, as mediated by Scp160), respectively, we determined whether Hhf1 and Scp160 work together or separately. To do this, we examined whether deletions of *HHF1* or *SCP160* in the separate mating partners or combined mutations in one of the mating partners had additive/synergistic effects upon mating. We crossed *hhf1Δ* cells against *scp160Δ* cells and observed a block in mating similar to that of *hhf1Δ* × WT crosses (Figure 5D), indicating that there was no additive effect when single gene mutations are present in the mating partners. However, when *hhf1Δ scp160Δ* double mutants are crossed against WT cells we observed a complete block in mating, reminiscent of a *scp160Δ* homozygous cross (Gelin-Licht et al., 2012). This indicates that Hhf1 and Scp160 contribute more or less equally to the mating process (in terms of quantitative mating efficiency) and act in an additive (non-epistatic) fashion.

### Histone H4 co-localizes with a mating pathway mRNA in the nucleus

Since histone H4 associates with mRNAs of the mating mRNP and is required for full RNP assembly and mating, we examined where it localizes in the cell. We found that RFP-tagged Hhf1 localized only to the nucleus and co-localized with nuclear (unexported) *STE2* mRNA (Figure 6A). We note that we could not detect any RFP-Hhf1 outside of the nucleus, where the bulk *of STE2* mRNA resides (*i.e*. on the ER) (Figure 6A). Thus, it would seem likely that the action of H4 on mRNP assembly occurs at the nuclear level.

**Figure 6.**
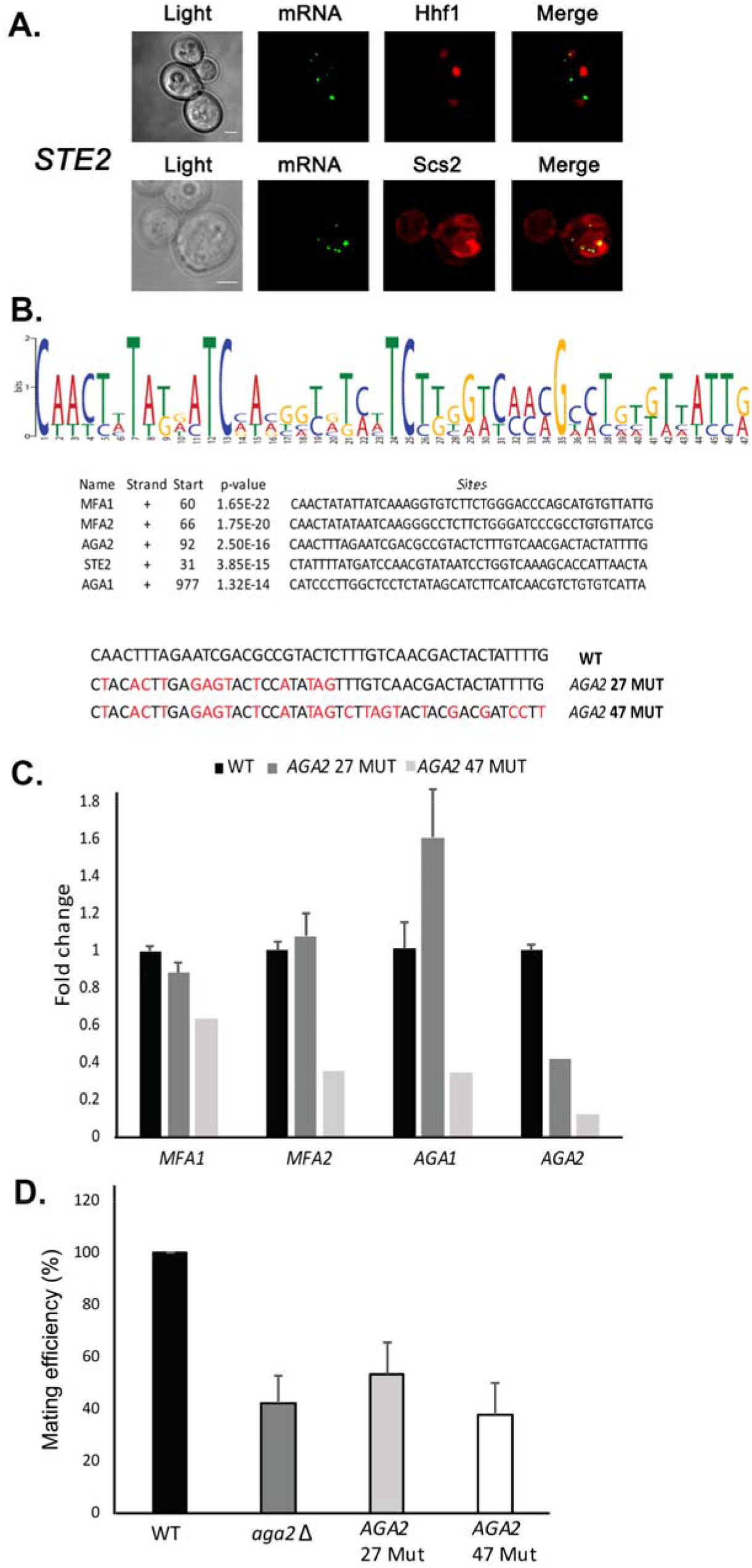
A motif in the mating mRNAs coding region facilitates mRNP assembly and mating. (A) Mating mRNAs and histone H4 co-localize only in the nucleus. Yeast expressing MS2 aptamer-tagged *STE2* mRNA from its genomic locus and MS2-CP-GFP(x3) were transformed with plasmids expressing either Hhf1-mCherry or Scs2-mCherry (an ER marker). *mRNA* - MS2-CP-GFP(x3) labeling; *Hhf1* – mCherry-Hhf1 labeling; *Scs2* – Scs2-mCherry labeling; *merge* – merger of mRNA and ER windows; *light* – transmitted light. Size bar = 2μm. (B) A conserved motif is common to all five *MAT*a mRNAs. MEME-ChIP analysis of the sequences of all five *MAT*a mRNAs was performed and revealed a 47 nucleotide consensus motif in the coding regions, shown schematically as a sequence logo based upon nucleotide representation. Also shown are two forms (*AGA2 27-MUT* and *AGA2 47-MUT*) in which the motif was altered within the coding region to remove conserved nucleotides without altering the amino acid sequence. (C) Mutation of the consensus motif in *AGA2* leads to an inhibition in mRNP particle assembly. MS2 aptamer-tagged *STE2* cells bearing the *aga2Δ* deletion were transformed with a single-copy plasmid expressing *AGA2* or either of the two *AGA2^mut^* forms. Next, WT, *AGA2 27-MUT*, and *AGA2 47-MUT* cells expressing MS2-tagged *STE2* were subjected to RaPID-qPCR with primers against *AGA1, AGA2* (note: not within mutated region), *MFA1*, and *MFA2*. The histogram indicates the levels of precipitated mating mRNAs from WT (black) or *AGA2 27-MUT* and *AGA2 47-MUT* (dark and light grey, respectively) cells after normalization for the level *of STE2* mRNA pulldown. (D) Deletion of *AGA2* or mutation of the consensus motif alters mating efficiency. *MAT*a wild-type cells (WT; BY4741), *AGA2 27-MUT or AGA2 47-MUT* mutant cells were crossed against wild-type *MAT*α cells and quantitative mating was assessed. Three biological repeats were performed; *p* <0.05.

### *MAT*a mating particle RNAs have a conserved sequence motif important for mRNA assembly, pheromone responsiveness, and mating

Since mating mRNAs multiplex with the help of histone H4, we examined their sequences for recognizable motifs using bioinformatic analysis (*e.g*. MEME Suite, MEME-ChIP) (Bailey et al., 2009). Although we anticipated that the H4 interaction site might reside (at least in part) within the 3’UTR, based upon the RaPID-MS experiment (Figure S1B), we identified a single motif of 47 nucleotides present in the coding region of all five *MAT*a messages examined (Figure 6B). To determine whether this motif facilitates RNA multiplexing and mating, we created two mutated versions in *AGA2* [*e.g*. a short 27 nucleotide version (*AGA2 27-MUT*) and a full-length 47 nucleotide version (*AGA2 47-MUT*)], which substituted certain conserved residues within the motif with synonymous mutations that did not alter the amino acid sequence. Using RNA-fold (http://rna.tbi.univie.ac.at/cgi-bin/RNAWebSuite/RNAfold.cgi), we found that the structure of *AGA2 47-MUT* was distorted, as compared to *AGA2 27-MUT* and the native *AGA2* motif (Figure S4A). When substituted for the native motif in *AGA2, we* found that *AGA2 47-MUT*, but not *AGA2 27-MUT*, led to a loss of *AGA2* mRNA assembly into the mating mRNP particle precipitated using *STE2* as the pulldown mRNA (Figure 6C). Similarly, the deletion of *AGA2* itself also led to defects in mating mRNP assembly (Figure S4B) and either motif mutation or gene deletion resulted in significant defects in the ability of cells to mate with wild-type yeast (Figure 6D). These results suggest that motif presence plays an important role in both mating mRNP particle formation and mating.

Bioinformatic analyses revealed that this motif is not present in the mating mRNAs expressed in *MAT*α cells, however, two distinct non-overlapping motifs of 45 and 25 bases could be identified in the *MF*α*1, MFA*α*2, STE3* and *SAG1* mRNAs (Figure S4C). Thus, both mating types appear to utilize different *cis* elements in their RNAs.

To determine whether particle assembly is ultimately important for both cellular responsiveness to the external pheromone signal (*i.e*. pheromone secreted by cells of the opposite mating type), and the secretion of pheromone and ability to induce a mating response in wild-type cells, we crossed *MAT*a cells expressing these *AGA2* mutants with GFP-labeled wild-type *MAT*α cells and assessed the shmooing efficiency of both cell types (Figure S5A). In parallel, we examined whether deletions in *HHF1, HHF2*, and *AGA1* also affected shmooing or the induction of shmoo formation in wild-type cells of the opposite mating type. We observed reduced shmoo formation for both the wild-type *MAT*a and *MAT*a deletion strains in the non-isogenic crosses, as compared to the isogenic WT *MAT*a and WT *MAT*a cross (Figure S5A). The reduction in shmoo formation (*i.e*. pheromone responsiveness) was detectable whether either histone H4 paralog was deleted or whether *AGA1* or *AGA2* were mutated. We note that *AGA2 27* mutant had the least effect upon shmooing, whereas the histone deletions were somewhat more effective.

These results suggest that defects in mRNP assembly affect not only the mutant cell’s ability to secrete pheromone and, thus, influence its wild-type mating partner, but also affect the mutant cell’s ability to respond to pheromone secreted by its wild-type mating partner. Therefore, mating mRNP particle assembly likely impacts upon both pheromone synthesis/secretion as well as pheromone responsiveness.

### P-body elements are not required for yeast cell mating

Our results show that mating mRNAs localize to nER in the cell body in both pheromone-treated and untreated cells (Figure 2C) and, in the case of *MFA2*, do not associate with P-bodies under conditions of endogenous gene expression (Haimovich et al., 2016). In contrast, it was proposed earlier that mating mRNAs, like *MFA2*, localize to P-bodies in the shmoo tip and that P-body components are necessary for transmission of the mating signal (Aronov et al., 2015). To re-examine the involvement of P-bodies in the mating process, we created deletions in *DHH1* and *PAT1* (*e.g. dhh1Δ, pat1Δ*, and *dhh1Δ pat1Δ*) and measured the ability of these mutants to mate with WT cells using the quantitative mating assay. However, unlike the previously published results (Aronov et al., 2015), we found that these same deletion mutants had no effect whatsoever upon mating (Figure S5B). Thus, we conclude that neither P-bodies nor P-body formation are necessary for transmission of the mating signal, nor are they *bona fide* sites for mating mRNA localization at endogenous levels of gene expression.

### Intergenic association between the *STE2* and *AGA2* genes

Since we observed *STE2* and *AGA2* mRNA colocalization both before and after nuclear export (Figures 2D and F), we hypothesized that their genes (located on ChrVI and ChrVII, respectively) might undergo interchromosomal interactions to facilitate RNA multiplexing upon transcription. To examine the possibility of allelic coupling, we performed chromatin conformation capture (3C) (Lieberman-Aiden et al., 2009), using multiple controls. First, we confirmed using qPCR that the chromatin was restriction enzyme-digested by >95 percent; second, we used a non-ligated control as a measure of background for ligation-dependent 3C signals; third, we used the *ASH1* gene as a negative control, as its mRNA does not associate with the mating mRNP particle; and fourth, to show allele-specific coupling we analyzed interactions between the *GET1* and *DDI2* genes, which are located in the same chromosomes as *STE2* and *AGA2*, respectively (Figure 7A and B). Importantly, we observed three interactions between *STE2* and *AGA2* consisting of ORF-ORF, ORF-5’UTR, and 3’UTR-3’UTR (Figure S6A), which were confirmed by DNA sequencing (Figure S6B). These interactions were specific, as they were not observed between *STE2* and *GET1, or AGA2* and *DDI2*, or between *ASH1* and *AGA2*, for example (see Figure S6A for example of latter).

**Figure 7.**
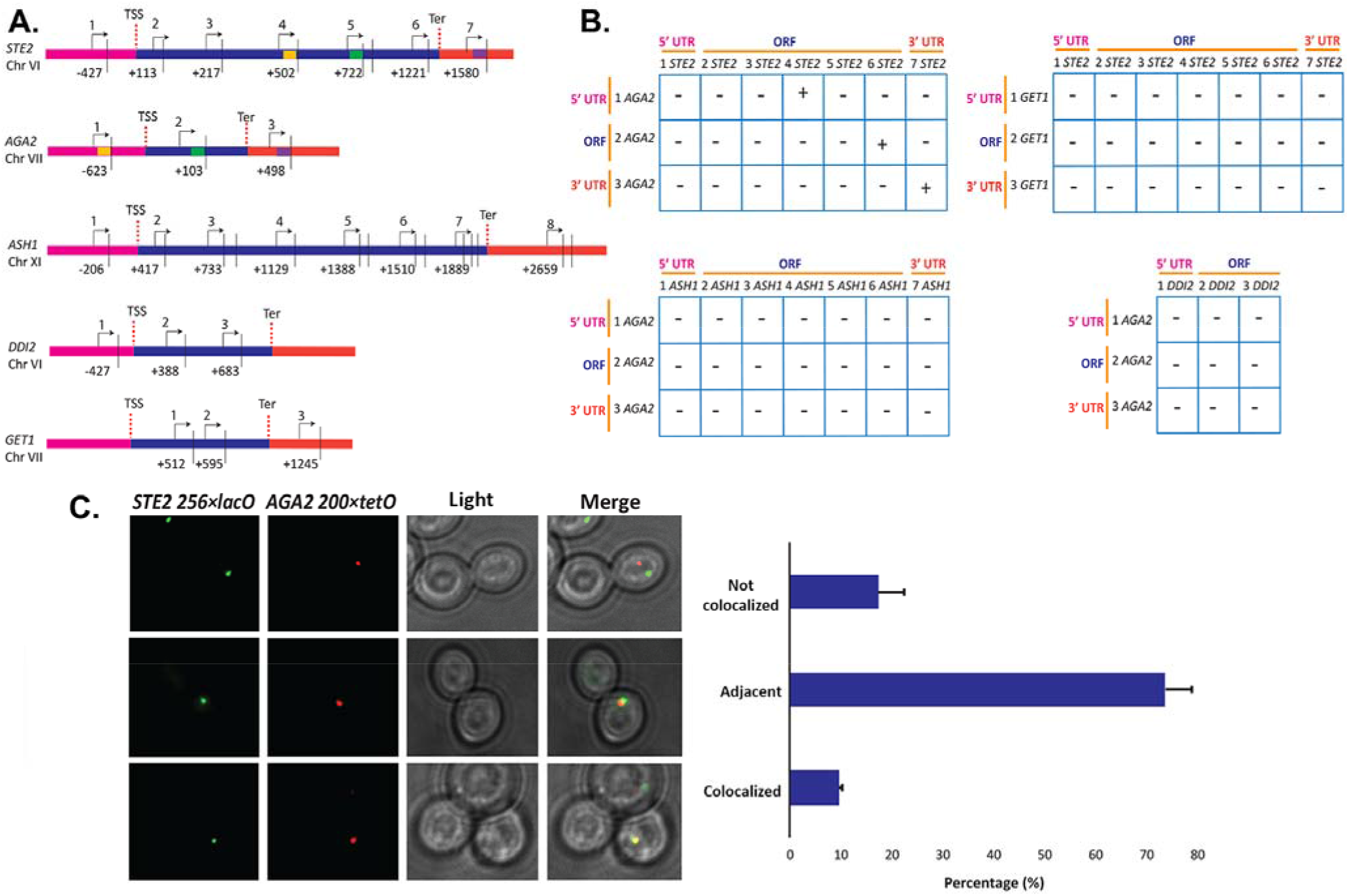
Intergenic association of *STE2* and *AGA2* genes. (A) Schematic of the *STE2, AGA2, ASH1, GET1 & DDI2* genes and the oligonucleotides used for their amplification from 3C DNA samples. Coordinates correspond to *Taq*I sites (shown as vertical black bars); site numbering is relative to ATG (+1). Forward primers used for 3C analysis were sense-strand identical (arrows) and positioned proximal to *Taq*I sites as indicated. Primers are numbered to distinguish the pairs used in the PCR reactions shown in *B*. 5’UTRs, ORFs and 3’UTRs are color-coded, as indicated. Transcription start sites (TSS) and termination sites (Ter) are indicated. (B) Matrix summarizing the intergenic association of represented genes as determined by 3C-PCR. Primer pairs corresponding to the different genes listed in *A* were used in PCR reactions. “+” indicates PCR amplification and the interaction between genes. “-” indicates no amplification. See Figure S6A for examples of both. 5’UTRs, ORFs and 3’UTRs are indicated by broken orange lines. (C) Live cell fluorescence microscopy of *AGA2-224×tetR* and *STE2-256×lacO* yeast grown to mid-log phase prior to imaging. Left panel: *STE2-256×lacO* gene is labeled with GFP-lacI; *AGA2-224×tetR* is labeled with tetR-tdtomato; *Merge* – merger *of STE2* and *AGA2* windows; *Light* – transmitted light. Size bar = 2μm. Right panel: histogram of the data from three biological replicas (avg.+std.dev.); *Co-localized* – fully overlapping signals; *adjacent* - partially overlapping signals; *Not co-localized* – no overlap between signals.

To confirm the interaction between the *STE2* and *AGA2* genes, we created yeast bearing *AGA2* tagged upstream of the 5’UTR with 224 tetracycline repressor repeats (224×tetR) and *STE2* tagged upstream of its 5’UTR with 256 lac operator repeats (256×lacO) and performed live fluorescence imaging using co-expressed tetR-tdTomato and GFP-tagged lad, respectively. We observed 9.5% fully overlapping signals and 73.6% partially overlapping signals co-localization, where both loci appear adjacent to each other (Figure 7C). In contrast, only 17.3% of the loci were not closely associated. As a control, we performed live imaging of *AGA2* 224×TetR and the *ASH1* gene tagged with 256×LacO repeats and the abovementioned fluorescent reporters (Figure S5C), and observed far less locus co-localization or adjacent loci (*e.g*. 2.3% and 27.2%, respectively), as compared to *AGA2* 224×TetR and *STE2* 256×LacO (Figure 7C). These results suggest that the *AGA2* and *STE2* loci exist in close association within the nucleus.

### mRNAs encoding heat shock proteins multiplex to form a heat-shock RNP particle

Chowdhary *et al*. (Chowdhary et al., 2019) first reported an intergenic association of heat shock protein (HSP) genes in yeast undergoing heat shock. If so, then we predicted that HSP mRNAs might also undergo multiplexing due to HSP gene allelic coupling. Thus, we tagged *HSP104* with the MS2 aptamer and performed RaPID to identify cohort HSP mRNAs in the pulldowns. Importantly, the mRNAs of HSP genes previously shown to undergo intergenic interaction (*e.g. TMA10, HSP12, HSP82, SSA2* and *SSA4*) (Chowdhary et al., 2019) were found to co-precipitate both before, but much more after heat shock (Figure 8A). To validate this result, we tagged *TMA10* with the PP7 aptamer in the cells expressing MS2-tagged *HSP104* and examined for mRNA co-localization upon expression of their respective fluorescent protein-tagged aptamer-binding proteins (*e.g*. PP7-PS-tomato and MS2-CP-GFP(x3)). We found that 59.5+7.7% (avg+std; n=3 experiments) of PP7-tagged *TMA10* mRNA granules co-localized with MS2-tagged *HSP104* mRNA (Figure 8B). Thus, HSP mRNAs appear to multiplex into an HSP mRNP particle during heat shock.

**Figure 8.**
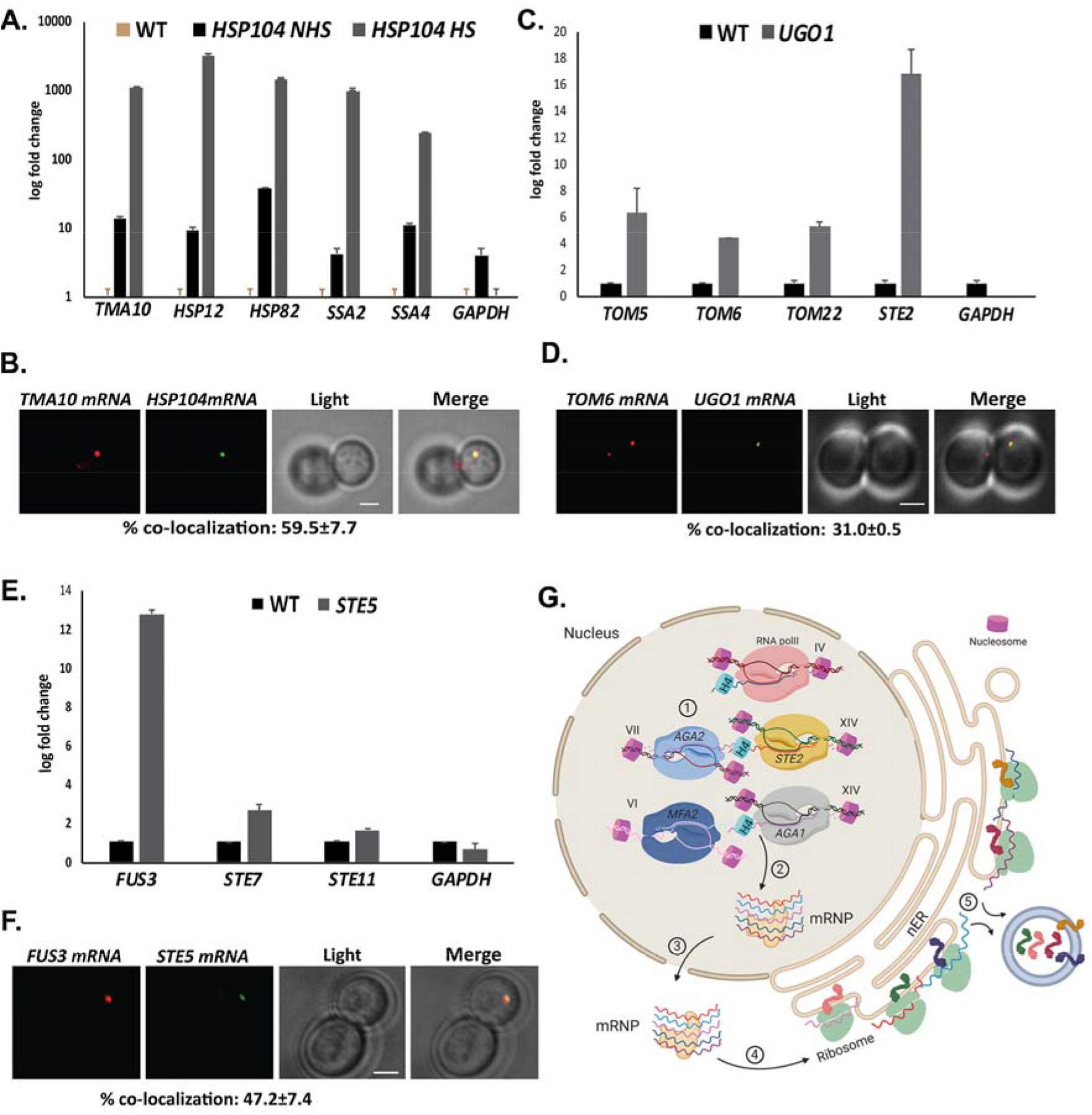
mRNAs encoding heat shock proteins, mitochondrial outer membrane proteins and MAPK protein multiplex to form different RNP particles. *MAT*a yeast strains (BY4741) expressing MS2 aptamer-tagged *HSP104, UGO1* or *STE5* from their genomic loci or untagged control (WT) cells were grown to mid-log phase (O.D._600_=0.5) and subjected to RaPID followed by qRT-PCR (RaPID-qRTPCR; see *Materials and Methods*). RNA derived from the total cell extracts or biotin-eluated fractions was analyzed by qRT-PCR using primer pairs corresponding to mRNAs expected to multiplex (see listed genes) or not (*e.g. GAPDH* mRNA). (A) mRNAs encoding HSPs multiplex upon heat shock. Cells expressing MS2 aptamer-tagged *HSP104* were either exposed to heat shock (10 min; 40°C; *HS*) or maintained at 30°C (*NHS*) prior to fixation and RaPID-qRT-PCR. (B) mRNAs of HSPs colocalize upon heatshock. Fixed cell fluorescence microscopy of yeast expressing MS2 aptamer-tagged *HSP104* and PP7 aptamer-tagged *TMA10* from their genomic loci, as visualized using MS2-CP-GFP(x3) and PP7-PS-tomato, respectively. Cells were exposed to heat shock prior to visualization. % co-localization (avg.+std.dev.) is given below, based on three biological replicas. *Merge* – merger of mRNA and nucleus windows; *light* – transmitted light. Size bar = 2μm. (C) MOMP mRNAs multiplex. Cells expressing MS2 aptamer-tagged *UGO1* were grown in medium containing glycerol as a carbon source, prior to fixation and RaPID-qRT-PCR. (D) MOMP mRNAs co-localize. Fixed cell fluorescence microscopy of yeast expressing MS2 aptamer-tagged *UGO1* and PP7 aptamer-tagged *TOM6* from their genomic loci, as visualized using MS2-CP-GFP(x3) and PP7-PS-tomato, respectively. (E) mRNAs encoding some MAPK pathway components multiplex. Cells expressing MS2 aptamer-tagged *STE5* were treated with α-factor (0.5μM) prior fixation and RaPID-qRT-PCR. (F) Fixed cell fluorescence microscopy of yeast expressing MS2 aptamer-tagged *STE5* and PP7 aptamer-tagged *FUS3* from their genomic loci, as visualized using MS2-CP-GFP(x3) and PP7-PS-tomato, respectively. (G) Model for mating mRNP particle formation and its role in the co-translation of secreted proteins involved in yeast cell mating. Yeast (*e.g. MAT*a cells, as shown) express mRNAs encoding soluble secreted and membranal proteins (*e.g*. Mfa1/Mfa2 and Aga1/Aga2/Ste3, respectively) from genomic loci located on different chromosomes (as listed) that undergo allelic coupling to form intergenic associations. The presence of histone H4 paralogs (Hhf1,2) allows the mRNAs assemble into a mating mRNA ribonucleoprotein (mRNP) particle that is exported to the cytoplasm and confers mRNA co-localization at the endoplasmic reticulum peripheral to the nucleus (nER). Co-translational translocation of the translated proteins at the ER may allow for coordination between the pathways responsible for the cellular responsiveness to pheromone of the opposite mating type (*i.e*. via Ste3 signaling and Aga1/Aga2 function in cell adhesion) and transmission of the cell’s own mating signal to cells of the opposite mating type (*i.e*. via Mfa1/Mfa2 secretion).

We also examined the HSP gene sequences for recognizable motifs and identified three potential motifs (Figure S7A) present in nearly all HSP cohort genes (Figure S7B). To determine whether histone H4 is involved in assembly of the HSP mRNP particle, we performed a growth test of the different histone deletion mutants upon heat shock. While single gene deletions did not affect doubling time after heat shock (Figure S7C), *AID-HHF1 hhf2Δ* cells treated with 3-IAA showed a longer doubling time after heat shock (at all times examined), as compared to WT cells or untreated *AID-HHF1 hhf2Δ* cells (Figure S7D). Thus, histone H4 appears to play a role in cell survival after heat-shock.

### mRNAs encoding mitochondrial outer membrane proteins or MAPK proteins also multiplex

To further identify examples of mRNA multiplexing, we employed existing MS2-tagged genes encoding proteins that undergo (or are likely to undergo) complex formation in yeast cells. We first examined whether mRNAs encoding mitochondrial outer membrane proteins (MOMPs) multiplex using tagged *UGO1* mRNA. Since the basal expression of *UGO1* mRNA was low (*i.e*. few puncta observed), we induced expression by growing cells on a non-fermentable carbon source (2% glycerol) before performing RaPID or subsequent fluorescence microscopy experiments. The pulldown of *UGO1* mRNA led to co-precipitation of cohort MOMP mRNAs (*e.g. TOM5, TOM6, TOM22*; Figure 8C), but not *GAPDH* mRNA, suggesting that a specific mRNA multiplex can be detected. Since we initially observed coprecipitation of the mating mRNAs along with *OM45* (Figure 1A and B), which codes for another MOMP, we examined for the precipitation of *STE2* mRNA with *UGO1* mRNA. Indeed, *STE2* mRNA could co-precipitate with *UGO1* mRNA, suggesting a possible interaction between MOMP and mating pathway mRNA complexes. To verify the interaction between MOMP mRNAs we tagged *TOM6* with the PP7 aptamer in cells expressing MS2-tagged *UGO1* and measured mRNA co-localization using MS2-CP-GFP(x3) and PP7-PS-Tomato, respectively. We found that 31.1+0.5% of PP7-tagged *TOM6* mRNA co-localized with MS2-tagged *UGO1* mRNA (Figure 8D). Thus, MOMP mRNAs appear to form multiplexes.

Finally, we examined whether mRNAs encoding non-secreted components of the yeast mating pathway (*e.g*. MAP kinase pathway genes: *STE5, FUS3, STE7*, and *STE11*) undergo multiplexing like mRNAs encoding their secreted counterparts (Figures 1, 2A and B, 3A). We tagged *STE5* mRNA with the MS2 aptamer and treated the cells for 1hr with α-factor (0.5μM) to induce gene expression. RaPID pulldown *of STE5* mRNA after pheromone induction led to the significant co-precipitation of *FUS3* mRNA, although only slight increases in *STE7* or *STE11* mRNA pulldown were detected (Figure 8E). We verified the association of MS2-tagged *FUS3* and PP7-tagged *STE5* mRNA in cells using MS2-CP-GFP(x3) and PP7-PS-Tomato, respectively, and observed 47.3+7.40% (avg+std;n=3 experiments) colocalization (Figure 8F). Thus, mRNAs for some components of the MAPK cascade may form multiplexes.

## Discussion

Prokaryotes coordinate the expression of genes encoding proteins involved in the same biological process/context by sequential placement in chromosomes in order to generate polycistronic messages (operons) that can be translationally controlled. Perhaps due to the greater complexity observed both at the gene and cellular levels, eukaryotes have preferentially relied on the discontiguous distribution of genes over different chromosomes. In this case, the co-regulation of gene expression presumably becomes dependent upon shared transcriptional control elements and specific RBPs that interact selectively with certain transcripts and confer trafficking to sites of co-translation. However, given that the number of RBPs is limited and that they can interact with a wide number of potential transcripts, it is unclear if and how defined subsets of mRNAs are packaged together in order to facilitate co-translation.

We hypothesized that mRNAs encoding proteins involved in the same process might assemble into macromolecular complexes composed of multiplexed transcripts and specific RBPs to form discrete RNP complexes. By performing mRNA pulldown and RNA-seq experiments we first identified a subset of mRNAs that encode secreted components involved in the mating of α-haploid cells. This mRNP complex included mRNAs encoding the a-pheromone receptor (*STE3)*, α-mating factor (*MF*α*1, MF*α*2*), α-agglutinin (*SAG1*), and a-pheromone blocker (*AFB1*) (Figures 1A and B, 2A, and S1). Correspondingly, we identified a mating mRNP particle from *MAT*a cells that included mRNAs encoding the α-mating factor receptor (STE2), a-mating pheromone (*MFA1, MFA2*), and a-agglutinins (*AGA1, AGA2*) (Figure 2B). Thus, we demonstrated that mRNAs coding for proteins of the same cellular process are able to multiplex into mRNP particles, presumably to allow for their delivery to the same intracellular site for local translation. Additional experiments in which we identified the existence of other mRNA subsets sets that multiplex, like those encoding HSPs, MOMPs, and MAPK pathway proteins (Figure 8) clearly support this idea. Thus, mRNA multiplexing *in trans* appears to form functional mRNPs or *transperons* [*i.e*. RNA operons; (Keene, 2007; Keene and Tenenbaum, 2002)] relevant to cell physiology (see below). We predict that analogous transperon particles containing different transcripts form within cells to modulate other physical processes. For example, recent work by Ashe and colleagues has shown that mRNAs encoding either translation factors or glycolytic enzymes co-localize and undergo co-translation in yeast (Morales-Polanco et al., 2020; Pizzinga et al., 2019). In addition, mRNAs encoding proteins in hetero-oligomeric/multisubunit complexes (proteasome, exocyst, COPI, FAS, ribosome, etc.) might form multiplexes to facilitate polypeptide complex assembly upon co-translation, as shown at the protein level (Shiber et al., 2018), even if dedicated assembly chaperones are needed. In our study, we may have missed multiplexes in the initial RaPID-seq experiment (Figures 1 and S1) for several reasons. First, RNA multiplexing may require elevated gene expression, as in the case of the HSP or MOMP complexes (Figure 8A-D), which require induction conditions to visualize the RNA granules, as opposed to the normal growth conditions employed initially. Second, the RaPID-seq pulldowns employed only 0.01% formaldehyde in the crosslinking procedure, as initially published (Slobodin and Gerst, 2010), and not the 0.1-0.5% formaldehyde we have employed in later work (Zabezhinsky et al., 2016). Third, the precipitated complexes were stringently washed and, thus, these different factors may have disrupted weaker transperons or missed them entirely.

Importantly, by using a variety of molecular and single-cell imaging techniques (*e.g*. 3C, live fluorescence imaging) we show specific and robust physical interactions between genes located on different chromosomes that encode mRNAs that form transperons (Figures 7, 8, and S6A and B). This observation of allelic coupling is strengthened by the work of Chowdhary *et al*. (Chowdhary et al., 2019), where the intergenic association of yeast HSP genes was first shown. Other chromatin conformation capture studies have revealed intergenic associations that lead to so-called transcription factories associated with different transcriptional regulators (*e.g*. promoters, transcription factors). For example, Schoenfelder *et al*. (Schoenfelder et al., 2010) showed interchromosomal interactions between Klf4-regulated globin genes in fetal liver cells that allowed for co-regulated gene transcription. Papantonis et *al*. (Papantonis et al., 2012) revealed that TNF-α responsive coding and miRNA genes in human endothelial cells undergo intrachromosomal interactions regulated by NFκB upon cytokine induction. Thus, specialized transcriptional factories created through allelic coupling may be a common mechanism for co-regulated gene expression in eukaryotes.

How allelic coupling comes about to form these factories and resulting transperons is not yet clear. However, we identified a role for histone H4 in the assembly of the mating transperon (Figures 3, 4, and S3) and its physiological consequences (*i.e*. pheromone responsiveness of both mating partners and mating) (Figures 4, S5A, S7D). RaPID-MS pulldowns of mating mRNAs encoding secreted proteins identified the H4 paralog, Hhf1, as binding to the *STE2, MFA1*, and *MFA2* mRNAs, but not to *ASH1* mRNA or *STE2* mRNA lacking its 3’UTR (Figure S2B). Thus, specific H4-RNA interactions are likely to be involved in transperon assembly. Hhf1 pulldowns directly precipitated the mating transperon (Figure 3A) and the deletion of either paralog (*HHF1* or *HHF2*) led to defects in RNA multiplexing (Figure 4C and D) and mating (Figures 4A and B, and 5D). Likewise, the combination of *HHF2* deletion and an auxin-induced degradative form of Hhf1 resulted in a complete block in mating under conditions where cell viability was not affected (Figures 4B and S3C). This suggests that Hhf1 and Hhf2 are important for the multiplexing of mating mRNAs encoding secreted components, and perhaps for the formation of other mRNA complexes identified. This idea is further borne out by experiments showing that histone H4 depletion also results in defects in cellular responsiveness to heat shock (Figure S7D).

Mutations in the amino terminal histone H4 acetylation sites had similar effects as gene deletions, whereby inactivating K-to-R mutations inhibited mating (Figure 5B and C). In contrast, K-to-Q mutations (or H4 overexpression) improved mating (Figure 5B). Thus, histone H4 paralogs, but no other histones (Figures 4A, S7C and D), appear to affect the transmission of physiological signals relevant with RNA multiplexing. More work is required to demonstrate the role histone H4 plays in transperon formation. Canonical histone functions relate mainly to DNA-protein interactions that facilitate nucleosome formation. However, the requirement of acetylation for H4 function in mating (Figure 5A-C) suggests that histone H4 might participate in the formation of a DNA-RNA intermediate upon gene transcription. This mechanism potentially allows for RNA multiplexing, provided that the loci encoding mRNAs intended for assembly *in trans* also are in close apposition, as evidenced here (Figures 7B, 8B and D, S6A and B).

Our lab is interested in how mRNA localization affects cell physiology and previous work already demonstrated that yeast mSMPs (including *MFA2*) localize primarily to nER (Kraut-Cohen et al., 2013). Here we verified that endogenously expressed *STE2* and *AGA1* mRNAs localize to nER, both before and after treatment with pheromone (Figure 2C), and that they co-localize both prior to and post nuclear export (Figure 2D-F). This demonstrates once again that mSMPs are not asymmetrically localized to the polarized extensions of yeast (*e.g*. bud and shmoo tips), unlike *ASH1* and polarity establishing mRNAs (*e.g. SRO7, SEC4, CDC42*) in budding cells (Aronov et al., 2007) or *SRO7* in shmooing cells (Gelin-Licht et al., 2012). It also reinforces the idea that multiplexing occurs post-transcriptionally within the nucleus and that translocated secreted and membrane proteins enter the nER and sort to the sites of secretion via the secretory pathway. The translocation of secreted and membrane proteins at the nER presumably imparts the temporal and spatial control required for the post-translational processing (*e.g*. glycosylation, proteolytic activation) needed for function, as opposed to entering the cER proximal to the bud or shmoo tips and adjacent to the site of secretion.

An earlier work (Aronov et al., 2015) suggested that mRNAs encoding mating components, like *MFA2*, localize preferentially to P-bodies in the shmoos of pheromone-treated yeast and that P-bodies are necessary for transmission of the mating signal and subsequent mating. However, these results were obtained under conditions of mRNA overexpression. In contrast, our previous work (Haimovich et al., 2016) found that endogenously expressed MS2-tagged *MFA2* mRNA does not localize to P-bodies, nor do P-bodies form under conditions of endogenous expression of the aptamer tagged mRNA. Here, we show that *MFA2* and other mating mRNAs (*e.g. MFA1, STE2*, and *AGA1*) do not localize to the shmoo tip, but rather to nER present in the cell body (Figures 2C and D, 6A, and S2A). Moreover, the deletion of genes necessary for P-body formation (*e.g. PAT1, DHH1*) had no effect upon mating (Figure S5B). Thus, it would appear that P-bodies are neither sites of localization for mating mRNAs, nor are necessary for transmission of the mating signal. These findings re-emphasize the importance of: (i) measuring mRNA localization under native conditions of gene expression; (ii) considering the possibility that RNA over-expression and/or the insertion of RNA aptamers may induce artifacts (*e.g*. mRNA mislocalization, P-body formation, or both); and (iii) examining mRNA integrity/localization using additional techniques (*e.g*. Northern analysis, RNA-seq, single-molecule FISH) (Garcia and Parker, 2015; Haimovich et al., 2016; Heinrich et al., 2017) for secondary validation.

Since *cis*-acting RNA elements may work in conjunction with specific RNA-binding proteins to confer multiplexing, we analyzed the coding regions and 3’UTRs of the mating mRNAs from the two yeast haplotypes. Sequence analysis revealed the presence of conserved motifs present in the mating mRNAs from either *MAT*a or *MAT*α cells (Figure 6B). We mutated the *MAT*a sequence element in the *AGA2* gene and found that it affected mRNP particle assembly similar to the deletion of *AGA2* (*aga2Δ*) (Figure 6C). Importantly, this correlated with an inhibition in cellular responsiveness (*i.e*. shmooing) towards its mating partner (Figure S5A), as well as the responsiveness of its mating partner (Figure S5A), resulting in significant defects in mating (Figure 6D). Thus, defects in transperon formation result in significant changes in cell physiology and we predict that a yet unidentified RBP interacts with this element to facilitate RNA assembly and, possibly, trafficking. The *MAT*a element is unlikely to be the H4-interacting motif itself, since it was not identified among the mating mRNAs from *MAT*α cells, which have other shared motifs (Figure S4C). Likewise, removal of the 3’UTR from *STE2* appeared to reduce H4 binding (Figure S2B), possibly indicating its role in histone association. Additional RaPID-MS experiments should help reveal the identity of this protein, as well as those interacting with the motifs in mating mRNAs from *MAT*α cells (Figure S4C), and genetic analyses should reveal their contribution to the mating process.

The results presented here support the concept that eukaryotic mRNP particles/transperons are the functional equivalents of bacterial operons, as first proposed by Keene and colleagues (Keene, 2007; Keene and Tenenbaum, 2002). Thus, eukaryotes may circumvent polycistronicity by assembling relevant mRNAs into functional complexes that confer co-translation. This is likely to occur via physical intergenic interactions that couple individual alleles concurrent with transcription. A schematic describing mRNA assembly into functional mating transperons via histone H4 mediation is shown in Figure 8G. While this work implies that there are higher order principles governing the selection of mRNAs for mRNP particle assembly, further study is required to uncover the entire mechanism by which mRNAs are recruited and assembled together into multiplexes, and to elucidate how histones and other proteins link mRNAs into fully formed transperons.

## Acknowledgements

The authors are grateful to Tsviya Olender (Weizmann Institute of Science) for bioinformatics help and Aviv Regev (Broad Institute, Cambridge, MA) for helpful support and for coining the term *transperon*. We also thank Amir Aharoni and Daniel Dovrat (Ben Gurion University, Beersheva), and Susan Gasser (Biozentrum, Basel) for strains and plasmids. This work was supported by grants to J.E.G. from the German-Israel Foundation (GIF; #I-1190-96.13/2012) and Minerva Foundation, Germany, as well as support from the Astrachan Olga Klein Fund (Weizmann Institute of Science) and the Israel Science Foundation (#578/18). C.N. and J.E.G. were partially supported by funding from NIH (NHGRI U54HG00306). R.R.N. was supported by a VATAT Fellowship for Postdoctoral Fellows from China and India. D.Z. is employed at Merck, Darmstadt, Germany. M.C.A.D. and H.S.S. study at the Free University, Berlin, Germany. H.S.S works at the Vitalant Research Institute, San Francisco, CA. J.E.G. holds the Besen-Brender Chair in Microbiology and Parasitology (Weizmann Institute of Science).

## Materials and Methods

### Yeast strains, genomic manipulations, growth conditions, and plasmids

Yeast strains used are listed in Table S1. Standard LiOAc-based protocols were employed for the introduction of plasmids and PCR products into yeast. For RNA and protein localization experiments, cells were inoculated and either grown at 26°C to mid-log phase (O.D._600_ = ~1) in standard rich growth medium containing 2% glucose (YPD) or in synthetic medium containing 2% glucose [*e.g*. synthetic complete (SC) and selective SC drop-out medium lacking an amino acid [25]. Induction of the methionine starvation-inducible plasmids (*e.g*. pUG36-based MS2-CP-GFP(x3)) was performed by transferring cells grown to mid-log phase to synthetic medium lacking methionine and subsequent growth of the cells for 1hr with shaking at 26°C. For growth tests, 2×10^7^ yeast cells grown to mid-log phase and normalized for absorbance at OD_600_ were serially diluted 1:10, plated by drops onto solid synthetic selective medium, and grown at different times/temperatures before photo-documentation. Three biological replicas were performed for each growth test.

Plasmids used are listed in Table S2. Gene deletions were made using the PFA6-based plasmids (Longtine et al., 1998) and PCR amplification of the deletion cassettes. Point mutations at genomic loci were made using a CRISPR/Cas9-based system (Mans et al., 2015) that was modified to express both Cas9 and gRNA from the same pRS426 plasmid. gDNAs were designed by CHOPCHOP software (Labun et al., 2016) and introduced between *SNR52* promoter and gRNA scaffold using the FastCloning technique (Li et al., 2011). The repair long primer was designed to have homologous overlapping sequences of 45 base pairs before and after the site of mutation, and was introduced into yeast along with Cas9 and gDNA in one transformation.

A *MAT*a W303 strain containing *GFP-lacI* at the *HIS3* locus and *tetR-3xCFP* at the *ADE1* locus (Dovrat et al., 2018) was modified by replacing *tetR-3xCFP* with *tetR-tdTomato*. The integration of *lacO* repeats and *tetR* repeats was performed as described in Dovrat *et al*. (2018) (Dovrat et al., 2018). A selection marker (*NAT*) was amplified with flanking sequences homologous downstream of the target gene (*e.g. STE2, AGA2, ASH1*) in the genome and transformed into yeast. After validation of correct insertion by PCR, a linearized plasmid containing either the *lacO* or *tetR* array (Dovrat et al., 2018), a selection marker (*e.g. LEU2* or *TRP1)*, and flanking *NAT* gene target sequences was transformed. This leads to replacement of the *NAT* selection marker by the linearized array. The integration of each array (*e.g. lacO* or *tetR*) into the target genes was performed sequentially and after visual confirmation of the first integration using fluorescence microscopy.

Growth assays to determine the doubling time of WT (BY4741) and histone mutants after heat shock involved growing the cells to mid-log phase at 30°C to O.D._600_ = 0.4-0.8. A portion of culture (0.5 O.D._600_ units) was maintained at 30°C and the remainder (0.5 O.D._600_ units) were subjected to heat shock at 50°C for 60 minutes (Figure S6C). Cells were then pelleted and resuspended in 1ml rich media (YPD; 24°C). A volume of 60 μl of cells was transferred to 120 μl of YPD media in 96-well plates and the O.D._600_ values were measured at at 30°C in 30 min intervals for 60 cycles using a Tecan liquid handling robot. For experiments involving heat shock and the auxin-mediated depletion of Hhf1 (Figure S6D), cells were grown to mid-log phase and auxin (3-IAA; 4mM final concentration) was added to the medium for 3.5 hrs at 30°C. After which, cells were exposed to heat shock at 50°C for 15, 30, or 60 min and then pelleted and resuspended in 1ml YPD and diluted and grown, as described above.

### Fluorescence microscopy

Cells grown for fluorescence microscopy were collected and fixed in a solution containing 4% (w/v) paraformaldehyde and 4% (w/v) sucrose for 20 min at room temperature, washed once, resuspended in 0.1M KPO4/1.2M sucrose (w/v) buffer solution, and kept at 4°C. For staining of mitochondria, Mitotracker Red (Molecular Probes, Eugene, OR, USA) was added to mid-log phase-grown cells to a final concentration of 0.2μM for 20min before harvesting. After collection, the cells were pelleted, washed once in fresh medium at the appropriate temperature, and analyzed by fluorescence microscopy. Representative images were acquired using either a Zeiss LSM780 or LSM710 confocal and a Plan-Apochromat 100x/1.40 oil objective (Carl Zeiss AG, Oberkochen, Germany). The following wavelengths were used (using separate tracks): LSM510 - GFP, excitation at 488nm/emission at >505nm; red fluorescence, excitation at 561nm/emission at >575nm; LSM710 – GFP, excitation at 480nm/emission at 530nm; monomeric RFP, excitation at 545nm/emission at 560-580nm. For statistical analyses, at least three identical individual experiments were carried out. Unless otherwise stated, a total of 100 RNA granules per experiment were scored for the mRNA localization in cells, while 100-150 cells per experiment were scored for protein localization. For the scoring of either RNA or protein localization, the average of n=3 experiments (± standard deviation) is given. For the P value, a student’s t-test (two-tailed, paired) was done.

### Single-molecule FISH

Yeast expressing MS2-tagged *STE2* and PP7-tagged *AGA2* were grown to mid-log phase. Cells were pelleted and suspended in PBS, fixed in the same medium upon the addition of paraformaldehyde (4% final concentration), and incubated at room temperature (RT) for 45min with rotation. Cells were washed three times with ice cold Buffer B (0.1M potassium phosphate buffer, pH 7.5 containing 1.2M sorbitol), after which cells were spheroplasted in 1ml of freshly prepared spheroplast buffer [Buffer B supplemented with 20mM ribonucleoside vanadyl complexes (Sigma-Aldrich, St. Louis, MO), 20mM β-mercaptoethanol, and lyticase (Sigma-Aldrich, St. Louis, MO) (25 U per O.D.600 unit of cells)] for 10min at 30°C. The spheroplasts were centrifuged for 5min at low speed (2500rpm; 660 x *g*) at 4°C and washed twice in ice cold Buffer B. Spheroplasts were then resuspended in 750ul of Buffer B and approximately 2.5 O.D._600_ units of cells were placed on poly-L-lysine coated coverslips in 12-well plates and incubated on ice for 30 min. Cells were carefully washed once with Buffer B, then incubated with 70% ethanol for several hours to overnight at −20°C. Afterwards, cells were washed once with SSCx2 (0.3M sodium chloride, 30mM sodium citrate), followed by incubating with Wash buffer (SSCx2 with 10% formamide), for 15 min at room temperature (RT; ~23°C). Next, 45μl of hybridization buffer (SSCx2, 10% dextran sulfate, 10% formamide, 2mM ribonucleoside vanadyl complexes, 1mg/ml *E. coli* tRNA, and 0.2mg/ml BSA) containing 50ng/μl MS2-Cy3 and PP7-Cy5 probe mix was placed on parafilm in a hybridization chamber. Coverslips with the immobilized cells were placed face down on top of the hybridization buffer and were incubated overnight at 37°C in the dark. After probe hybridization, cells were incubated twice in Wash buffer for 15min at 37°C. Next, the cells were washed once with SSCx2 containing 0.1% Triton X-100, incubated with SSCx2 supplemented with 0.5μg/ml DAPI for 1min at RT and finally washed with SSCx2 for 5min at RT. Cells were mounted with Prolong Glass (Thermo Scientific) mounting media on clean microscope slides. Samples were imaged using a Zeiss AxioObserver Z1 DuoLink dual camera imaging system equipped with an Illuminator HXP 120 V light source, PlanApo 100× 1.4 NA oil immersion objective, and Hamamatsu Flash 4 sCMOS cameras. Incremental (0.2μm) z-stack images were taken using a motorized XYZ scanning stage 130×100 PIEZO and ZEN2 software at 0.0645μm/pixel. Images were processed by deconvolution. At least 50 cells with mRNA spots, were scored using the FISHquant program (Mueller et al., 2013) (https://bitbucket.org/muellerflorian/fish_quant) to analyze deconvolved images of single cells. mRNA co-localization was scored manually and was defined by overlap between the deconvolved signals. Three biological replicas were performed for each experiment and the average ± standard deviation is given.

### RaPID procedure for precipitation of RNP complexes and RNA sequencing

The pulldown of MS2 aptamer-tagged mRNAs and detection for bound RNAs and proteins was performed using the RaPID procedure, essentially as described (Slobodin and Gerst, 2010; Slobodin and Gerst, 2011). Yeast strains bearing endogenously expressed genes tagged with the MS2 aptamer (Haim et al 2007) were grown in a volume of 400ml to midlog phase at 26°C with constant shaking to an O.D._600_ = ~0.8. Cells were centrifuged in a Sorvall SLA3000 rotor at 1100 x *g* for 5min, resuspended in 200ml of complete synthetic medium lacking methionine in order to induce the expression of the MS2-CP-GFP-SBP protein, and grown for an additional 1hr. The cells were collected by centrifugation as described above, washed with PBS buffer (lacking Ca^++^ and Mg^++^), and transferred into a 50ml tube and pelleted as above. For the experiment shown in Figure 1, proteins were cross-linked by the addition of 8ml PBS containing 0.01% formaldehyde and incubated at 24°C for 10min with slow shaking. For all other experiments, 0.05% formaldehyde was used. The cross-linking reaction was terminated by adding 1M glycine buffer pH=8.0 to a final concentration of 0.125M and additional shaking for 2min. The cells were then washed once with ice-cold PBS buffer and the pellet was flash frozen in liquid nitrogen, and stored at −80°C.

For cell lysis and RNA pulldown, cell pellets were thawed upon the addition of icecold lysis buffer (20mM Tris-HCl at pH7.5, 150mM NaCl, 1.8mM MgCl2, and 0.5% NP40 supplemented with Aprotinin [10 mg/ml], PMSF [1mM], Pepstatin A [10mg/ml], Leupeptin [10 mg/ml], 1mM DTT, and 80 U/ml RNAsin [Promega]) at 1mL per 100 O.D._600_ U, and 0.5ml aliquots were then transferred to separate microcentrifuge tubes containing an equal volume of 0.5 mm glass beads, and vortexed in an Vortex Genie Cell Disruptor (Scientific Instruments, New York, USA) shaker at maximum speed for 10min at 4°C. Glass beads and unbroken cells were sedimented at 4°C by centrifugation at 1700 x *g* for 1 min, and the supernatant removed to new microcentrifuge tubes and centrifuged at 11000 *×g* at 4°C for 10min. The total cell lysate (TCL) was then removed to a fresh tube and protein concentration was determined using the microBCA protein determination kit (Pierce). Protein (10mg of total cell lysate) was taken per pulldown reaction. In order to block endogenous biotinylated moieties, protein aliquots were incubated with 10μg of free avidin (Sigma) per 1mg of input protein for 1h at 4°C with constant rotation. In parallel, streptavidin-conjugated beads (Streptavidin-sepharose high performance, GE Healthcare) were aliquoted to microcentrifuge tubes according to 5μl of slurry per 1mg of protein (not >30μl overall), washed twice with 1ml of PBS, once with 1ml of lysis buffer, and blocked with a 1:1 mixture of 1ml of lysis buffer containing yeast tRNA (Sigma; 0.1mg/100ml of beads) and 1ml of 4% BSA in PBS at 4°C for 1 h with constant rotation. Following blocking, beads were washed twice in 1ml of lysis buffer. Pulldown was performed by adding the indicated amount of avidin-blocked TCL to the beads, followed by incubation at 4°C for 2–15h with constant rotation. Yeast tRNA was added to the pull-down reaction (0.1mg/tube) to reduce nonspecific interactions. We used standard 1.7ml microcentrifuge tubes when working with small volumes of TCL or 15ml sterile polypropylene centrifuge tubes with larger volumes. Following pull-down, the beads were centrifuged at 660 x *g* at 4°C for 2min; the supernatant then removed, and the beads washed three times with lysis buffer (*e.g*. 1ml volume washes for small tubes, 2ml for large tubes), twice with wash buffer (20mM Tris pH7.5, 300mM NaCl, and 0.5% Triton X-100), all performed at 4°C (each step lasting 10min with rotation). The beads were then equilibrated by a final wash in 1-2ml of cold PBS, pelleted by centrifugation as above, and excess buffer aspirated. For elution of the crosslinked RNP complexes from the beads, 150ml of PBS containing 6mM free biotin (Sigma) was added to the beads, followed by 1h of incubation at 4°C with rotation. After centrifugation at 660 x *g* for 2min, the eluate was transferred into a fresh microcentrifuge tube, re-centrifuged, and transferred into another tube to assure that no beads were carried over. To reverse the cross-link, the eluate was incubated at 70°C for 1-2h with an equal volume of 2X cross-link reversal buffer (100mM Tris pH7.0, 10 mM EDTA, 20mM DTT, and 2% SDS) for RNA analysis or with an appropriate volume of 5X protein sample buffer (5X: 0.4 M Tris at pH 6.8, 50% glycerol, 10% SDS, 0.5 M DTT, and 0.25% bromophenol blue) for protein analysis using SDS-PAGE. For RaPID pulldowns and RT-PCR analysis three biological replicas were performed each and the average ± standard deviation is given.

For the RNA sequencing and analysis shown in Figures 1 and S1, libraries were constructed for RNA-Seq from the affinity purified RNA, as described for the control (nonstrand-specific) library in (Levin et al., 2010), except that the oligo(dT) selection and RNA fragmentation procedures were omitted. In addition, only random hexamers were used for cDNA synthesis and Ampure beads (Beckman-Coulter, IN, USA) were used for size selection. Libraries were sequenced using an Illumina HiSeq2000 sequencer (paired-end 76 base reads) to a depth of ~10^7^ reads. Gene expression levels were estimated from the RNA-Seq data using Kallisto (Bray et al., 2016) and targeting the *S. cerevisiae* reference transcriptome (derived from the Saccharomyces Genome Database and leveraging the *Saccharomyces cerevisiae* S288C genome version R64-2-1), where we added 100 bases to the 3’ UTR of each coding region. Differentially expressed genes were identified using edgeR (Robinson et al., 2010), with the dispersion parameter manually set to 0.1. Those genes reported as at least 4-fold differentially expressed (FDR<0.001) were retained as significantly differentially expressed. After log transforming the data (log2(TPM+1)), the gene expression values from the control sample were subtracted from experimental samples and plotted in a heatmap illustrating expression levels as compared to the control sample. Expression quantitation, differential expression, and plotting of heatmaps were facilitated through use of the transcriptome analysis modules integrated into the Trinity software suite (Haas et al., 2013). This experiment was performed twice and gave similar results – a representative experiment is shown in Figures 1 and S1, and Table S1.

For protein analysis, samples of the eluate from were electrophoresed on 10% SDS-PAGE gels, followed by silver staining using the Pierce Silver Stain kit (ThermoFisher, USA) to identify bands of interest. Bands were excised from gels and processed for reversed-phase nano-liquid chromatography-electrospray ionization tandem mass spectrometry at the Israel National Center for Personalized Medicine (Weizmann Institute, Rehovot, Israel).

### Immunoprecipitation

The immunoprecipitation of protein-RNA complexes was as described in Gelin-Licht *et al*. (Gelin-Licht et al., 2012). Briefly, WT (BY4741) cells, WT cells expressing HA-tagged Hhf1, and WT cells expressing both MS2 aptamer-tagged *STE2* and HA-Hhf1 were grown to midlog phase (O.D._600_ = ~0.5) in 200ml synthetic selective medium at 26°C. Cultures were centrifuged at 3,000 x *g* for 5min at 4°C and resuspended in 1ml of lysis buffer (20mM Tris-Cl pH 7.5, 150mM KCl, 5mM MgCl2, 100U/ml RNasin, 0.1% NP-40, 1mM dithiothreitol, 2μg/ml aprotinin, 1μg/ml pepstatin, 0.5μg/ml leupeptin, and 0.01μg/ml benzamidine). Cells (100 O.D._600_ units each) were broken by vortexing with glass beads (0.5mm size) for 10min at 4°C and centrifuged at 10,000 × *g* for 10 min at 4°C to yield the total cell lysate (TCL). For IP, 3mg of TCL was diluted in lysis buffer (final volume, 1 ml) and subjected to IP with 3μl of monoclonal anti-HA antibody (16B12; Covance) overnight at 4°C with rotation. Next, 50μl of a protein G agarose slurry (Santa Cruz Biotechnology, Dallas, TX, USA) was added and the samples were incubated for an additional 1h at 4°C with rotation. The precipitates were washed (×5) with lysis buffer and protein–RNA complexes were eluted by incubation at 65°C for 30 min in 300μl of elution buffer (50mM Tris-HCl pH 8.0, 100mM NaCl, 10mM EDTA, 0.1% SDS, and 100μg/ml proteinase K in RNasefree water).

For RNA detection, RNA was extracted from each sample using the Epicentre MasterPure kit (Lucigen, WI, USA), treated with DNase (2hrs), and 15.1μl (from the total of 35μl) were taken for reverse transcription (RT). For RT-PCR, 1μl of the RT reaction was diluted (x4) and 1μl was taken for PCR detection using specific primers for *STE2, MFA1, MFA2*, as well as *EXO70, ASH1, SRO7*, and *UBC6*, as potential controls. For qRT-PCR, 1μl of the RT reaction was diluted (x4) and 1μl taken for qRT-PCR detection using SYBR Green (as described above) with specific primers for *AGA1, AGA2, STE2, MFA1, MFA2*, and *UBC6*.

For Western analysis, samples (30μg) were electrophoresed on 15% SDS-PAGE gels. Protein detection using anti-HA (Covance 16B12, BioLegend, Dedham, MA, USA) or antiactin antibodies (Cell Signaling, Danvers, MA, USA) was performed using the Amersham ECL Western Blotting Detection Kit (GE Healthcare Life Sciences). Quantification of protein bands in gels was performed using GelQuant.NET software provided by biochemlabsolutions.com.

### Quantitative Mating Assay

Quantitative analysis of yeast mating was performed based on the complementation of auxotrophic markers present in the *MAT*a and *MAT*α cells after crossing (*i.e*. the *met15* mutation in BY4741 *MAT*a cells and *lys2* mutation in BY4742 *MAT*α cells are complemented upon mating to yield diploids that grow on medium lacking methionine and lysine). Cells from each mating type were grown to mid-log phase at 26°C to O.D._600_ = 0.3-0.4. 10^6^ cells from each mating type (in 150μl volume), as well as 1:1 *MAT*a and *MAT*α mixtures, were collected by filtration on individual cellulose nitrate filters (25mm diameter, 0.45μm pores; BA-85 Schleicher & Schuell, Maidstone, UK) placed on the top of a working filtration surface of a disposable 500ml filtration apparatus (Corning, NY, USA). After filtration to remove the liquid medium, each filter was placed cell-side up on YPD plates for 3hrs at 30°C, prior to resuspension in 10ml of SC-Met,-Lys in a sterile 50ml tube, followed by vortexing, serial dilution (1:10,100,1000) and plating of 100μl of the dilutions onto SC-Met,-Lys plates to select for diploids. Mating efficiency was determined by colony scoring after 3 days at 26°C. Three biological replicas were performed for each experiment and the average ± standard deviation is given.

### Quantitative yeast pheromone responsiveness (shmoo formation)

To determine the responsiveness of cells towards their mating partner (*e.g*. WT *MAT*α cells towards mutant *MAT*a cells and vice versa), the level of shmoo formation in each strain was measured. Cells from each mating type were grown to mid-log phase at 26°C to O.D._600_ = 1-2. Next, 10^6^ cells from each mating type were mixed (1:1) in a final volume of 150μl and collected by filtration on individual cellulose nitrate filters (25mm diameter, 0.45μm pores; BA-85 Schleicher & Schuell, Maidstone, UK) placed on the working filtration surface of a disposable 500ml filtration apparatus (Corning, NY, USA). After filtration to remove the liquid medium, each filter was placed cell-side up on YPD plates for 3hrs at 30°C, prior to resuspension in 1ml of 1X TE, followed by vortexing and pelleting of the cells. Cells were then fixed for 15min by resuspension in 4% (w/v) paraformaldehyde and 4% (w/v) sucrose for 20 min at room temperature, washed once, suspended in 0.1M KPO4/1.2M sucrose (w/v) buffer solution, and kept at 4°C. Images were acquired using Zeiss Plan-Apochromat 100x/1.40 oil objective (Carl Zeiss AG, Oberkochen, Germany). For statistical analyses, at least three identical individual experiments were carried out. Unless otherwise stated, a total of 100 or more cells per strain per experiment were scored for the formation of shmoo projections. For scoring of shmooing efficiency, the average of n=3 experiments is given. For the P value, a one way Anova with multiple comparison was performed.

### Chromosome Conformation Capture

The quantitative chromosome conformation capture method, TaqI-3C (Chowdhary et al., 2019; Hagege et al., 2007), was employed with modifications for use in yeast. BY4741 *MAT*a cells were cultured to 0.6-0.8 O.D._600_ in a volume of 50 ml to early log phase at 30°C and then crosslinked with 1% formaldehyde. Crosslinked cells were then subjected to glass bead-mediated lysis in FA lysis buffer (50 mM HEPES pH 7.9, 140 mM NaCl, 1% Triton X-100, 0.1% sodium deoxycholate, 1 mM EDTA, 1 mM PMSF) for two cycles (20min each) of vortexing at 4°C. Cell lysates were collected as described according to the RaPID procedure (*i.e*. beads and unbroken cells were sedimented at 4°C by centrifugation at 1700 x *g* for 1 min, and the supernatant removed to new microcentrifuge tubes and centrifuged at 11,000 x *g* at 4°C for 10min). After centrifugation, a thin translucent layer of chromatin was observed on the top of the pellet of cell debris. The supernatant was discarded and the pellet resuspended in 1 ml FA lysis buffer. The resuspended material was centrifuged at 13,000 x *g* for 10 min at 4°C and the resulting pellet resuspended in 500 μl of 1.2X *Taq*I restriction enzyme buffer. Next, 7.5 μl of 20% (w/v) SDS (final concentration of 0.3% SDS) was added and the sample incubated for 1 h at 37 °C with shaking (900 rpm). Then 50 μl of 20% (v/v) Triton X-100 (final concentration = 1.8%) was added and the sample further incubated for 1 h at 37 °C with shaking (900 rpm). An undigested sample of genomic DNA (50 μl aliquot) was removed and stored at −20 °C to determine enzyme digestion efficiency. To the remaining sample, 200 U of *Taq*I (New England Biolabs) was added and incubated at 60°C overnight. The next day, 150 μl of digested sample was heat-inactivated at 80°C for 20 min in the presence of added SDS (24 μl 10% SDS; final concentration = 1.7%). To samples of digested material (174ul each), 626ul of 1.15X ligation buffer and 80ul of 10% Triton X-100 (final concentration of 1%) were added and incubated at 37^0^ for 1 hour, while shaking gently. Proximity ligation in the sample was performed using 100 U of added T4 DNA ligase (New England Biolabs) at 4°C for 16h. The ligated samples were then digested with RNase (final concentration of 11 ng/μl; RNaseA and 28 U/ μl RNAseT1; ThermoScientific) at 37°C for 20 min. Proteinase K (final concentration of 56 ng/μl; Sigma Aldrich) digestion was performed at 65°C for 12 h (note: final concentration of SDS= 0.4%). The 3C DNA template was extracted twice using an equal volume of phenol-chloroform (1:1 TE-saturated phenolchloroform v/v), followed by extraction with an equal volume of phenol-chloroform-isoamyl alchohol (25:24:1 v/v), followed by extraction with an equal volume of chloroform-isoamyl alcohol (24:1 v/v). The DNA was precipitated in the presence of 2ul of glycogen (20mg/ml), sodium acetate (0.3 M final concentration, pH5.2), and 2.5 volumes of ethanol at room temperature overnight. The 3C DNA template obtained was stored at −20°C. DNA concentration was determined by absorption spectroscopy at 260 nm. Typically, 125–500 ng of the 3C DNA template was used in the PCR reactions. The PCR product was further eluted and sequenced for confirmation.

## Supplemental Figure Legends

**Figure S1.**
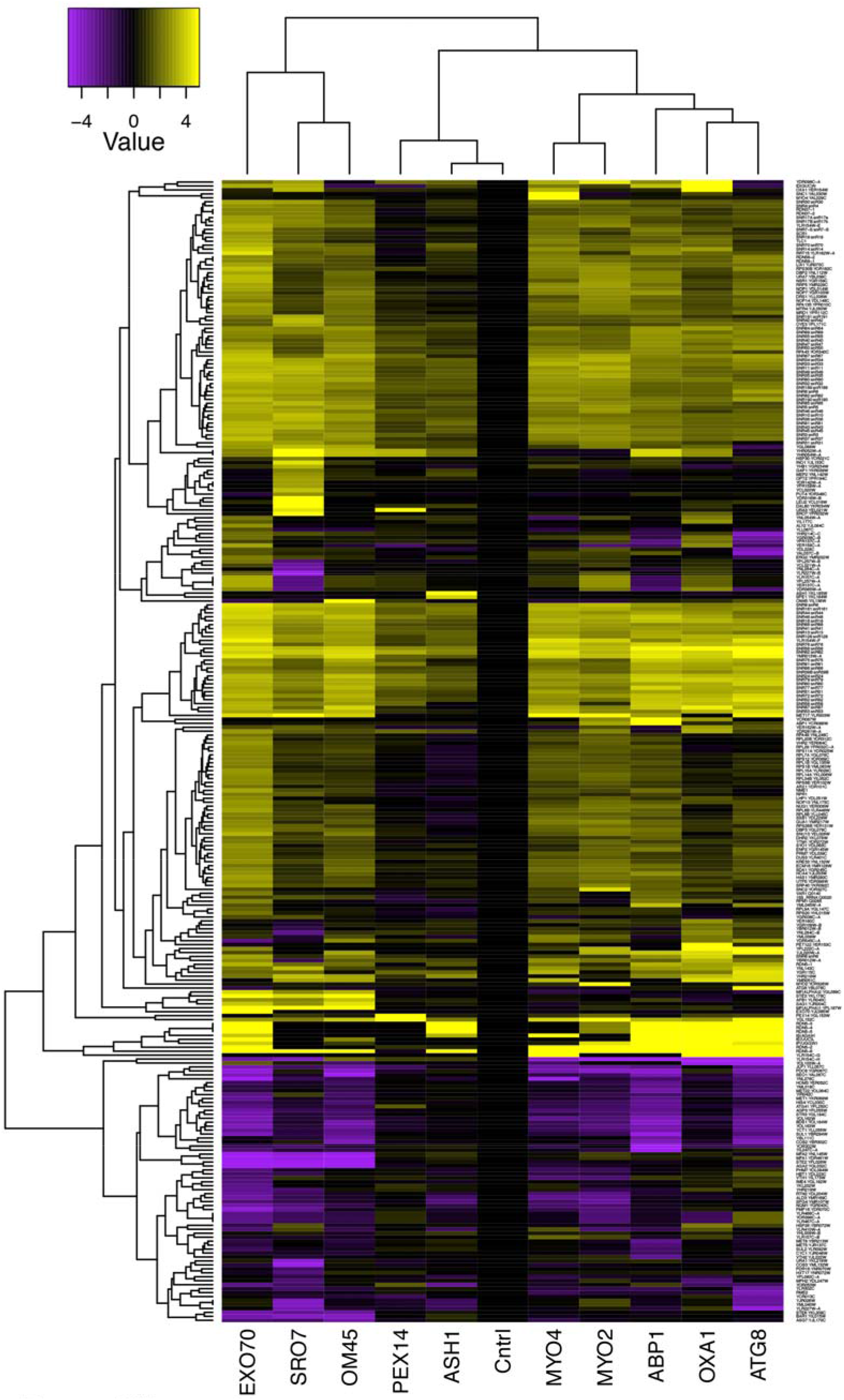
mRNAs encoding secreted and membrane proteins involved in yeast mating multiplex into a single ribonucleoprotein particle. An expanded version of the figure shown in Figure 1A is given. Different MS2 aptamer-tagged mRNAs (listed on x axis) expressed from their genomic loci were precipitated from *MAT*α yeast (BY4742) by MS2-CP-GFP-SBP after formaldehyde crosslinking *in vivo* and cell lysis procedures (RaPID; see Materials and Methods). RNA-seq was performed and the reads (including those of non-coding RNAs, including Ty elements, tRNAs, snoRNAs, ribosomal RNAs, etc.) were plotted to yield a heatmap for the relative enrichment of the tagged as well as non-tagged mRNAs (see color key for approximate values). Red-hatched rectangles indicate the target tagged mRNAs.

**Figure S2.**
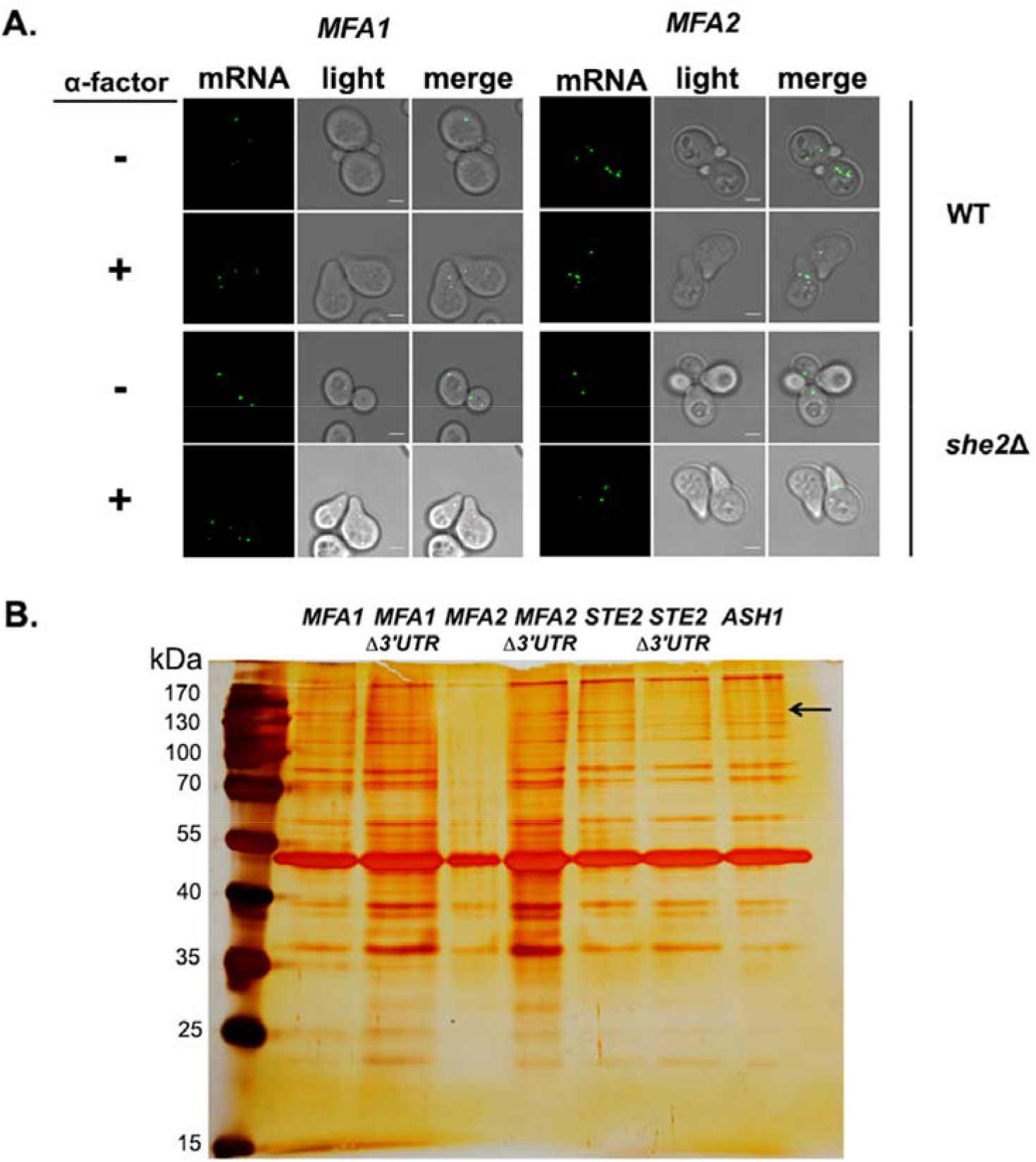
*MFA1* and *MFA2* mRNAs localize to the cell body and identification of a RNA-binding protein that binds to the 3’UTR of the mating mRNAs. (A) *MFA1* and *MFA2* mRNAs localize to the cell body with or without pheromone treatment, and in the presence or absence of She2. *MAT*a wild-type and *she2Δ* yeast expressing MS2-CP-GFP(x3) from a singlecopy plasmid and either MS2 aptamer-tagged *MFA1* or *MFA2* from their genomic loci were grown to mid-log phase on selective medium, either treated with or without added pheromone (10μM; 90min), and scored for RNA localization by confocal microscopy. (B) Identification of an RNA-binding protein (Hhf1) that binds to the 3’UTR of *STE2* mRNA. Yeast strains expressing MS2 aptamer-tagged *MFA1, MFA2*, and *STE2* either with or without their 3’UTRs, and *ASH1* were grown and processed for RaPID-MS (see Materials and Methods). The eluate after biotin-release was resolved by SDS-PAGE on a 10% polyacrylamide gel and silver stained. The arrow indicates the band of ~150kDa that exists in the *STE2* lane (and all *MFA1* and *MFA2* lanes), but not in the *ASH1* orSTE2Δ3’UTR lanes. *kDa* indicates kilodaltons.

**Figure S3.**
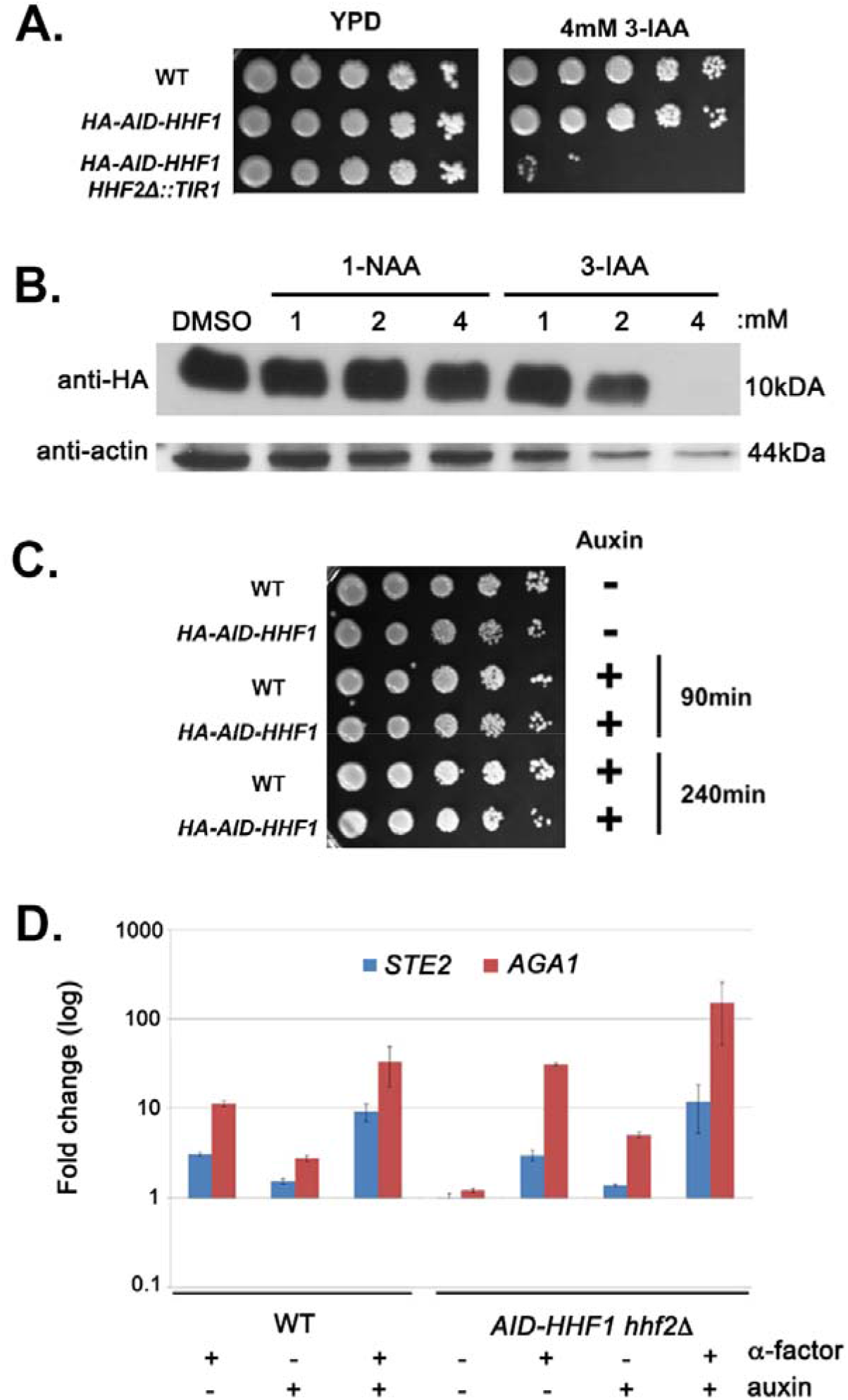
Conditional removal of both histone H4 paralogs leads to synthetic lethality overextended time. (A) Long-term treatment of *HA-AID-HHF1 hhf2Δ* cells with auxin leads to lethality. Wild-type (WT; BY4741) yeast and *HA-AID-HHF1* strains either bearing or lacking *TIR1* integration into the *HHF2* locus (*i.e*. the *hhf2Δ* mutation) were grown on liquid YPD to mid-log phase (O.D._600_=0.5). One O.D._600_ unit from each strain was removed, washed, and diluted serially 5 times (10-fold each). Cells were spotted onto pre-warmed solid YPD medium either with or without 4mM of 3-IAA, and grown for 48hrs at 26°C prior to photodocumentation. (B) HA-AID-Hhf1 disappears upon treatment with 4mM 3-indole acetic acid. *HA-AID-HHF1 hhf2Δ* cells were grown to mid-log phase prior to treatment with DMSO or different concentrations of 1-NAA or 3-IAA (auxin) for 2hrs. After harvesting and total cell lysate (TCL) preparation, samples (30μg) were resolved on 15% SDS-PAGE gels, transferred to nitrocellulose membranes, and probed with antibodies against HA and actin. (C). *HA-AID-HHF1 hhf2Δ* cells remain viable for at least 4hrs of treatment with 4mM 3-IAA. The indicated wild-type (BY4741) and *HA-AID-HHF1 hhf2Δ* cells were grown on rich medium containing glucose (YPD) to mid-log phase (O.D._600_=0.5) and treated for the indicated times with 4mM 3-IAA. One O.D._600_ unit from each strain was removed, washed, and diluted serially 5 times (10-fold each). Cells were spotted onto the same medium and grown for 48hrs at 26°C, prior to photodocumentation. (D) Neither auxin treatment nor histone H4 depletion abolishes pheromone-induced gene expression. WT and *HA-AID-HHF1 hhf2Δ* cells were grown to mid-log phase prior to treating cells with either auxin (3-IAA; 4mM), pheromone (10uM α-factor), or both for 2hrs. After treatment, total cell lysates were prepared, RNA extracted, and analyzed by qRT-PCR using primers against *STE2* and *AGA1*.

**Figure S4.**
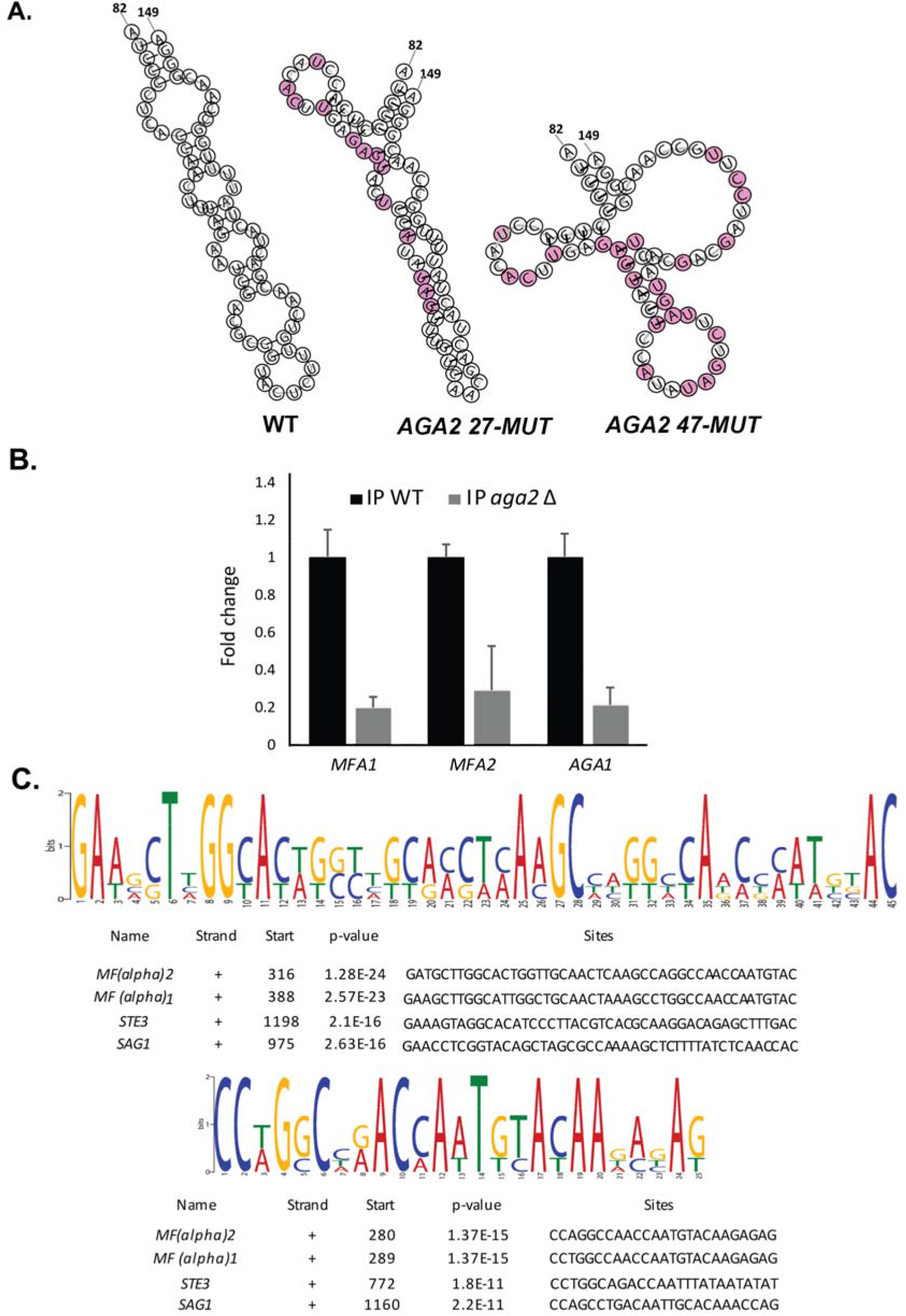
Structure of native and mutated *AGA2* mating motifs; effect of *AGA2* deletion on mating mRNP assembly; and identification of consensus motifs in *MAT*α mating mRNAs. (A) *AGA2 47-MUT* has a distorted structure. RNA-fold was used to model the native and mutant versions of the mating mRNA consensus motif in *AGA2*. Mutations in *AGA2 27-MUT* and *AGA2 47-MUT* are illustrated in pink. (B) Deletion of *AGA2* reduces the multiplexing of mating mRNAs. Wild-type (WT) *MAT*a (BY4741) and *aga2Δ* mutant yeast expressing MS2 aptamer-tagged *STE2* from its genomic locus were grown to mid-log phase (O.D._600_=0.5) and subjected to RaPID followed by RT-PCR (RaPID-PCR; *see Materials and Methods*). RNA derived from the total cell extract or the biotin-eluated fraction was analyzed by qPCR with primers against *AGA1, MFA1*, and *MFA2*. The histogram indicates the levels of precipitated mating mRNAs from either WT (black) and *aga2Δ* (grey) cells after normalization for *STE2* pulldown. (C) Two consensus motifs are common to the four *MAT*α mating RNAs. MEME-ChIP analysis of the sequences of all four *MAT*α mating mRNAs was performed and revealed two conserved motifs (sites) of 45 (upper sequence logo) and 25 (lower sequence logo) nucleotides, as shown schematically for each gene. *p* values are listed; “+” = sense strand.

**Figure S5.**
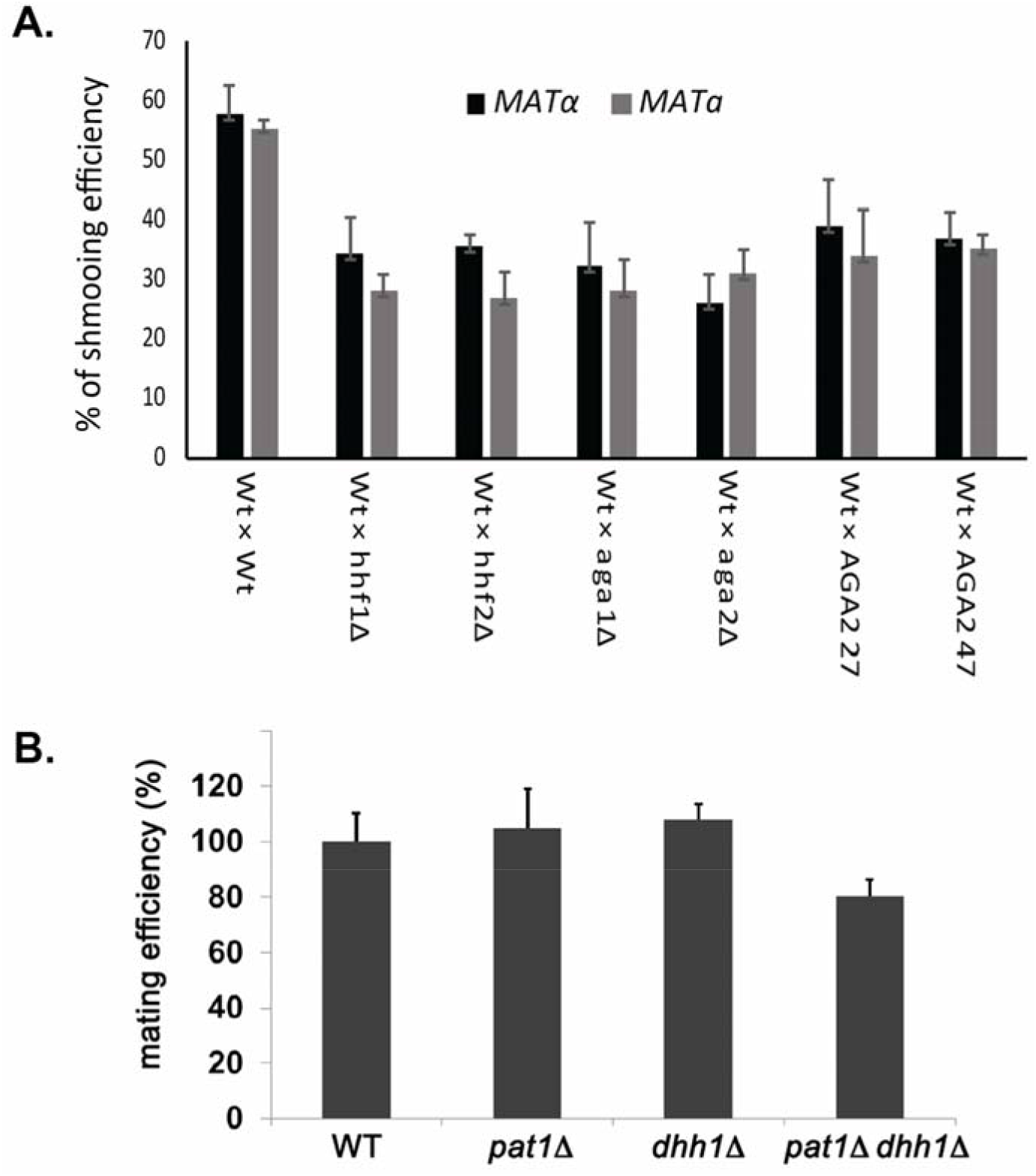
Mutations in histone H4 or *AGA2* in *MAT*α cells alters both mating partners responsiveness to pheromone; P-body components are not required for the mating response. (A) Mutations in the histone H4 paralogs or *AGA2* alter *MAT*α cell responsiveness to pheromone, as well as in their WT *MAT*a mating partner. *MAT*a wild-type (WT; BY4741), *hhf1Δ, hhf2Δ, aga1Δ, aga2Δ*, and *AGA2^mut^* cells were crossed against WT *MAT*α cells and quantitative mating was assessed. Later cells were fixed in 4% paraformaldehyde in medium containing 3.5% sucrose, and analyzed using widefield microscope to determine the percentage of shmooing (*i.e*. pheromone-responsive) cells. Three biological repeats were performed and gave similar results. (B) P-body components are not required for efficient mating. *MAT*a wild-type (WT; BY4741) cells and the *pat1Δ, dhh1Δ*, and *pat1Δ dhh1Δ* deletion mutants were crossed against WT *MAT*α cells and quantitative mating was assessed. Three biological repeats were performed and gave similar results; *p* <0.05.

**Figure S6.**
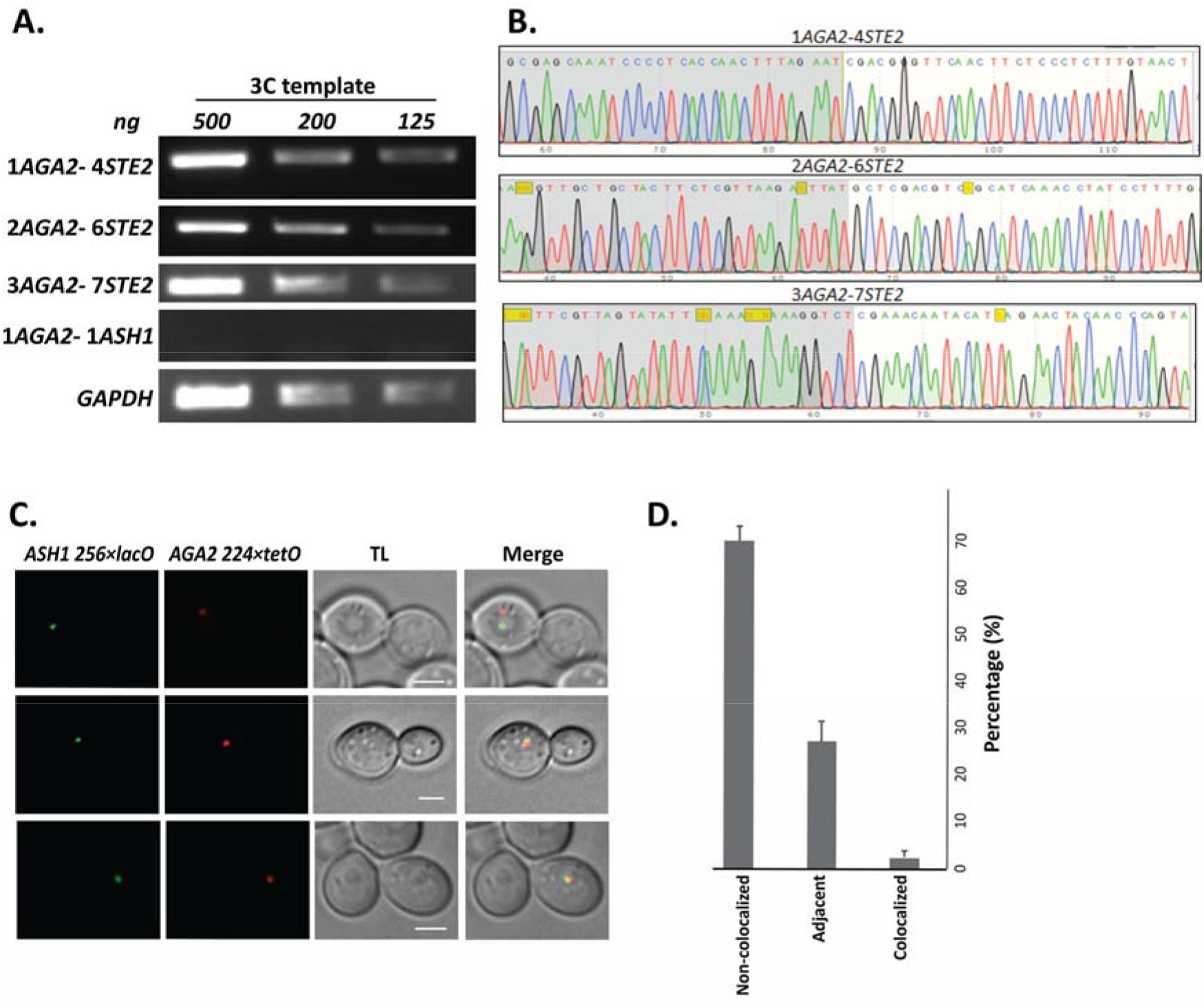
Intergenic association of *STE2* and *AGA2* genes. (A) *STE2* and *AGA2* genes undergo allelic interactions. PCR products derived reactions using the indicated primer pairs (to the genes shown in Figure 8A) and 3C-processed DNA were fractionated on agarose gels (1.5%) and visualized by ethidium bromide staining using an AlphaImager 2000. Lanes represent the 3C-PCR output using the indicated concentrations of DNA template (ng DNA). *GAPDH* primers was used as a control for the PCR reaction. (B) Sequence confirmation of the *STE2-AGA2* gene interaction. Positive PCR products were eluted from the gel and sequenced to confirm the nature of the observed band. A DNA sequencing chromatogram representing the observed sequence of the different *STE2* and *AGA2* chimeric PCR products is shown. (C) *AGA2* and *ASH1* genes do not undergo allelic interactions. Live cell fluorescence microscopy of yeast having the *AGA2* and *ASH1* genes tagged with *224×tetR* and *256×lacO* repeats, respectively. Cells were grown to mid-log phase prior to imaging. *ASH1* 256×LacO repeats were visualized by GFP-lacI; AGA2-224×TetR was visualized by tetR-td tomato; *merge* – merger of *STE2* and *AGA2* windows; *TL* – transmitted light. Size bar = 2μm. Right panel: histogram of the data from three biological replicas (avg.+std.dev.); *Co-localized* – fully overlapping signals; *adjacent* - partially overlapping signals; *Not co-localized* – no overlap between signals.

**Figure S7.**
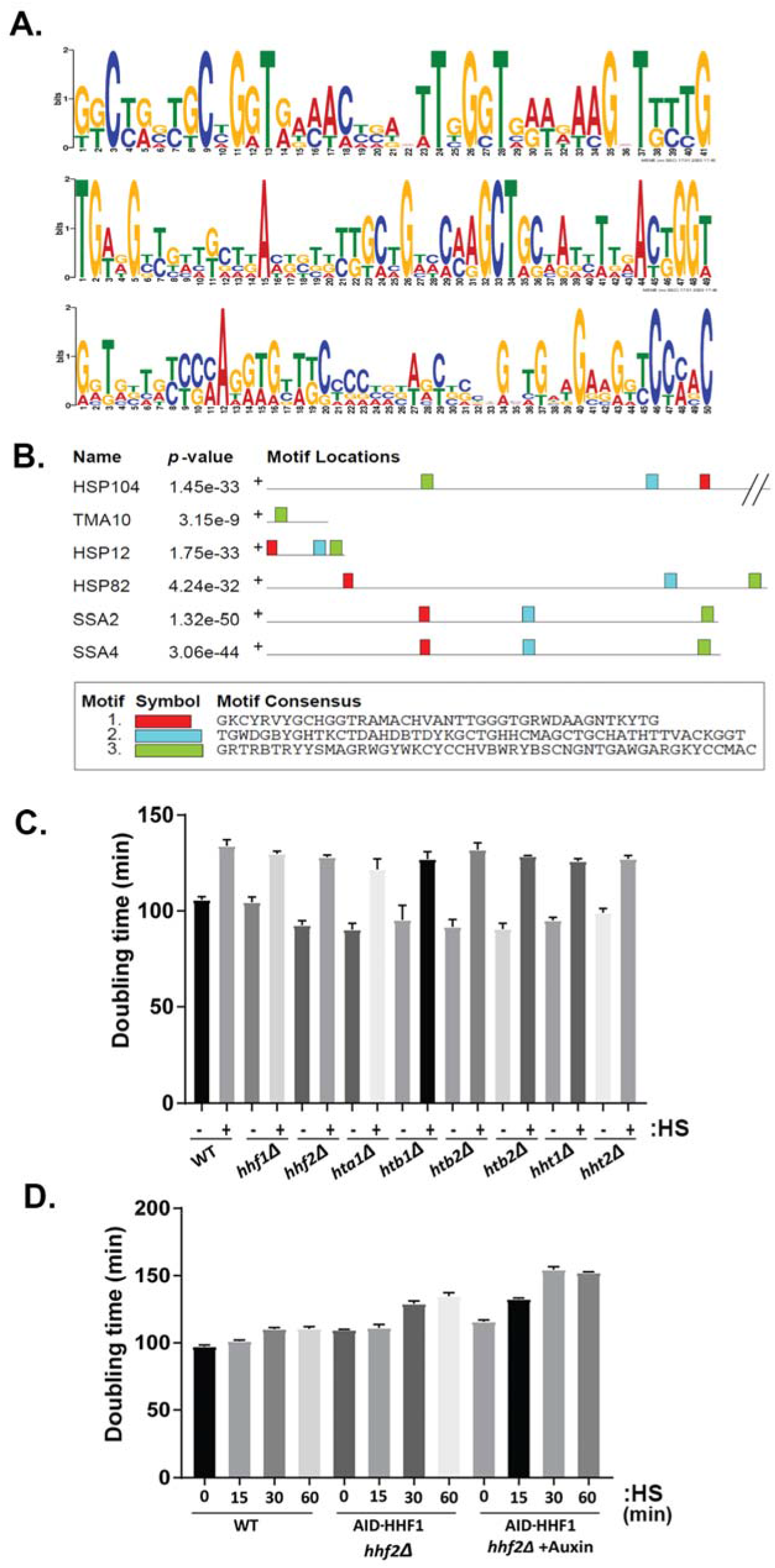
Motifs in HSP mRNA coding regions and role of histone H4 in RNP formation. (A) A conserved motif is common to all six HSPs mRNAs. MEME-ChIP analysis of the sequences of all six mRNAs was performed and revealed three conserved motifs in the coding regions, shown schematically as sequence logo-based upon nucleotide representation. (B) Distribution of the motifs over the HSP genes. Motif location is plotted, along with their *p* values over the different genes. Motifs 1-3 correspond to those shown in A, respectively. (C) Single gene histone deletions do not affect cell growth after heat shock. The doubling times (minutes) of WT, *hhf1Δ, hhf2Δ, hta1Δ, hta2Δ, htb1Δ, htb2Δ, hht1Δ, and hht2Δ* strains were assessed for cells grown on liquid rich medium (YPD) exposed to heat shock (50°C; 60min; +*HS*) or not (-*HS*) and then allowed to recover for 18 hrs at 30°C. Error bars represent the avg.+std.err.mean) of four biological repeats. (D) Combined *hhf2Δ* and *AID-HHF1* alleles result in increased doubling time upon auxin induction and heat shock. *MAT*a wild-type (WT) and *hhf2Δ AID-HHF1* cells were grown to mid-log phase, treated either with or without auxin (3-IAA; 4mM) for 3.5 hrs, and exposed to heat shock (50°C; 0, 15, 30, 60min; *HS*). After removal to fresh YPD medium lacking auxin, strains were grown for 18 hrs at 30°C and the doubling times assessed. n = 3 experiments; *p* value <0.05.

## Supplementary Tables

**Table S1. RNA-sequencing Data (Excel table)**

Different MS2 aptamer-tagged mRNAs (listed on column heads) expressed from their genomic loci or control cells (Cntrl) lacking a tagged mRNA were precipitated from *MAT*α yeast (BY4742) by MS2-CP-GFP-SBP after formaldehyde crosslinking *in vivo* and cell lysis procedures (RaPID; see Materials and Methods). RNA-seq was performed, the results for each gene are listed on the following sheets. Sheet 1 (Counts.matrix) represents the number of reads assigned to each gene by Kallisto. Sheet 2 (edgeR_DE_results) represents all genes identified by edgeR, as being differentially expressed in the corresponding pairwise comparison to the control, requiring a false discovery rate (FDR) ≤ 0.1. Sheet 3 (DE_expr_sub_cntrl) represents gene length-normalized expression values, log2 transformed followed by subtraction of the control expression value. Sheet 4 (Expr_targets_sub_cntrl) represents the Sheet 3 values restricted to the baits (target mRNAs).

**Table S2.**
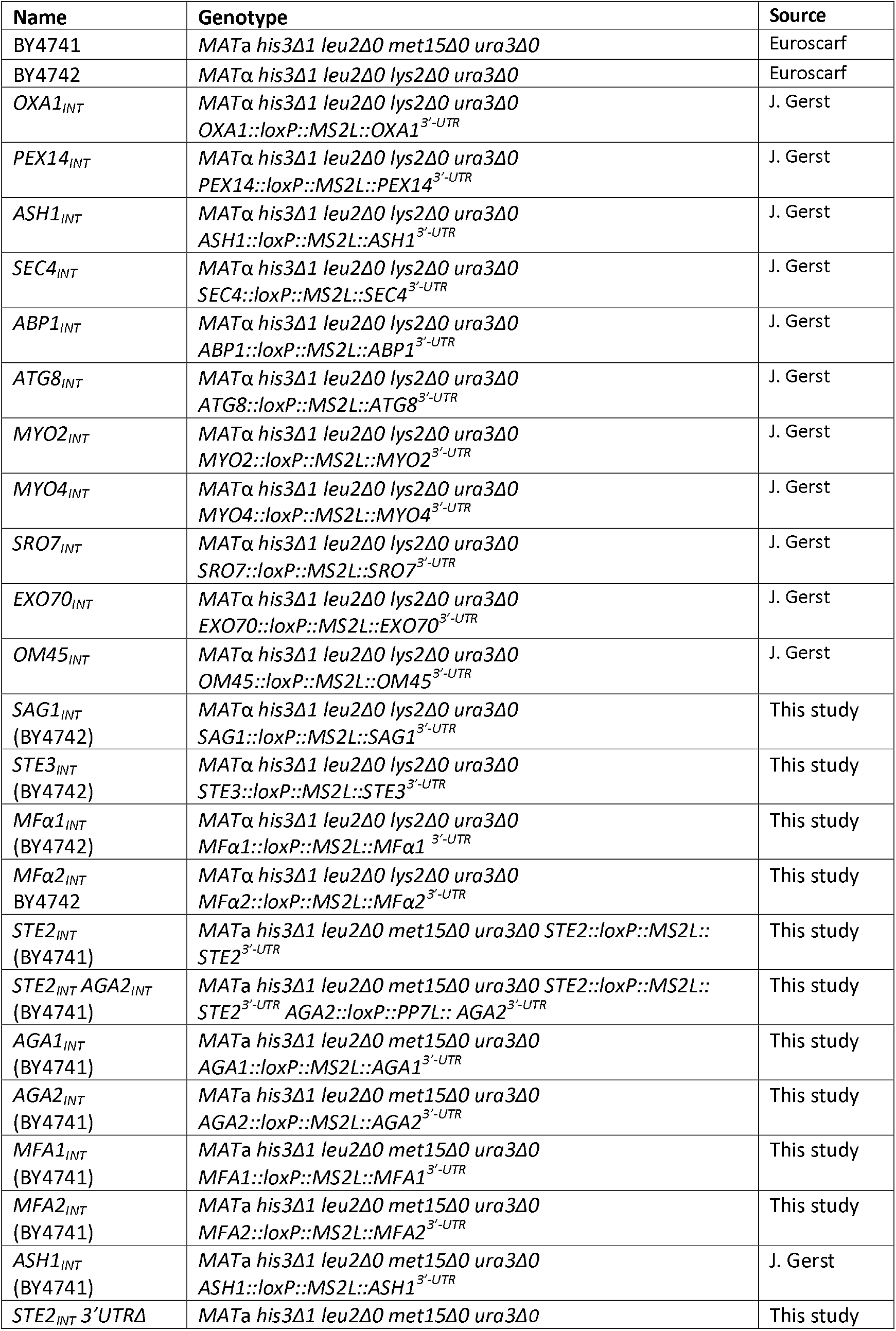
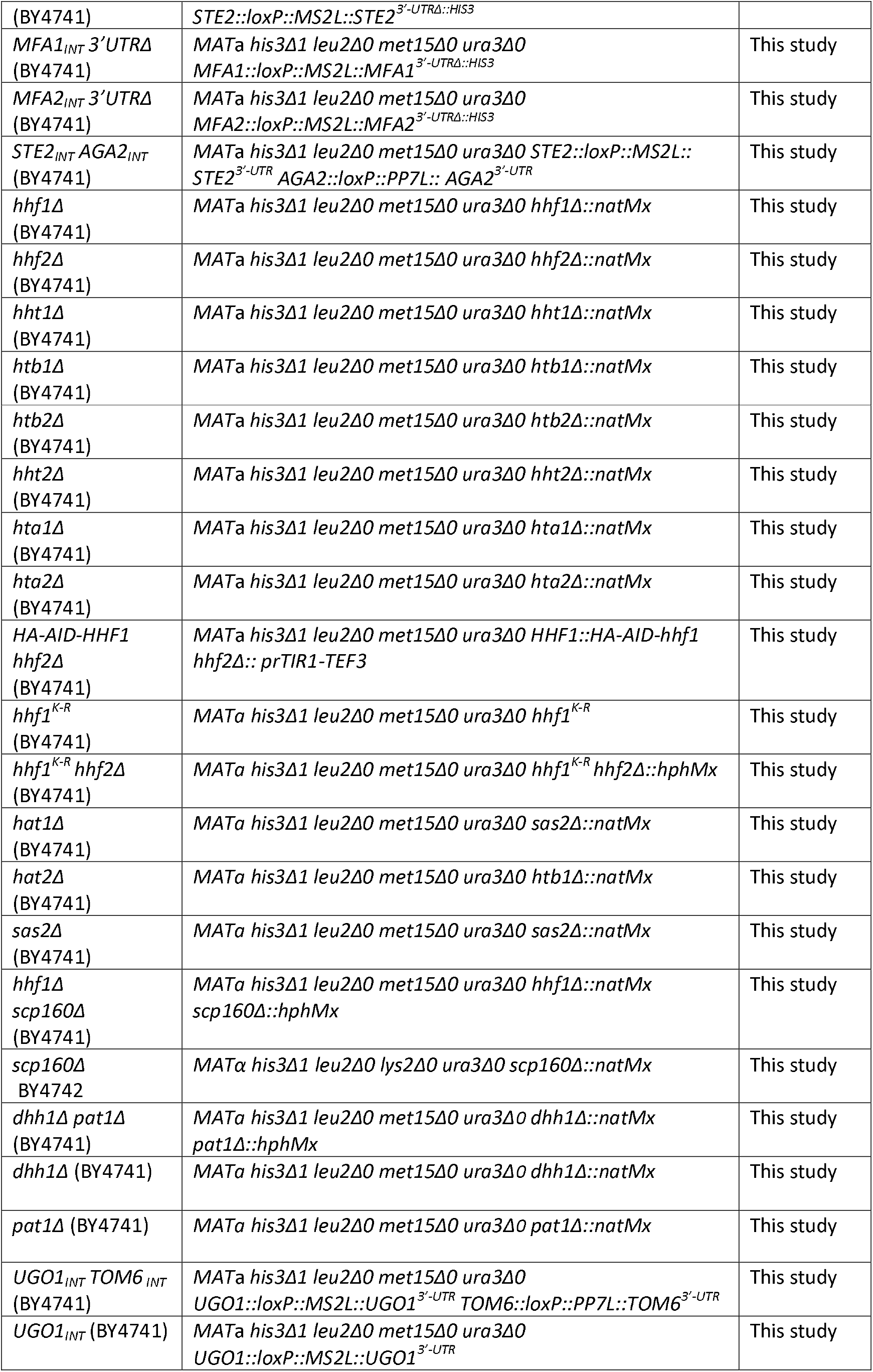
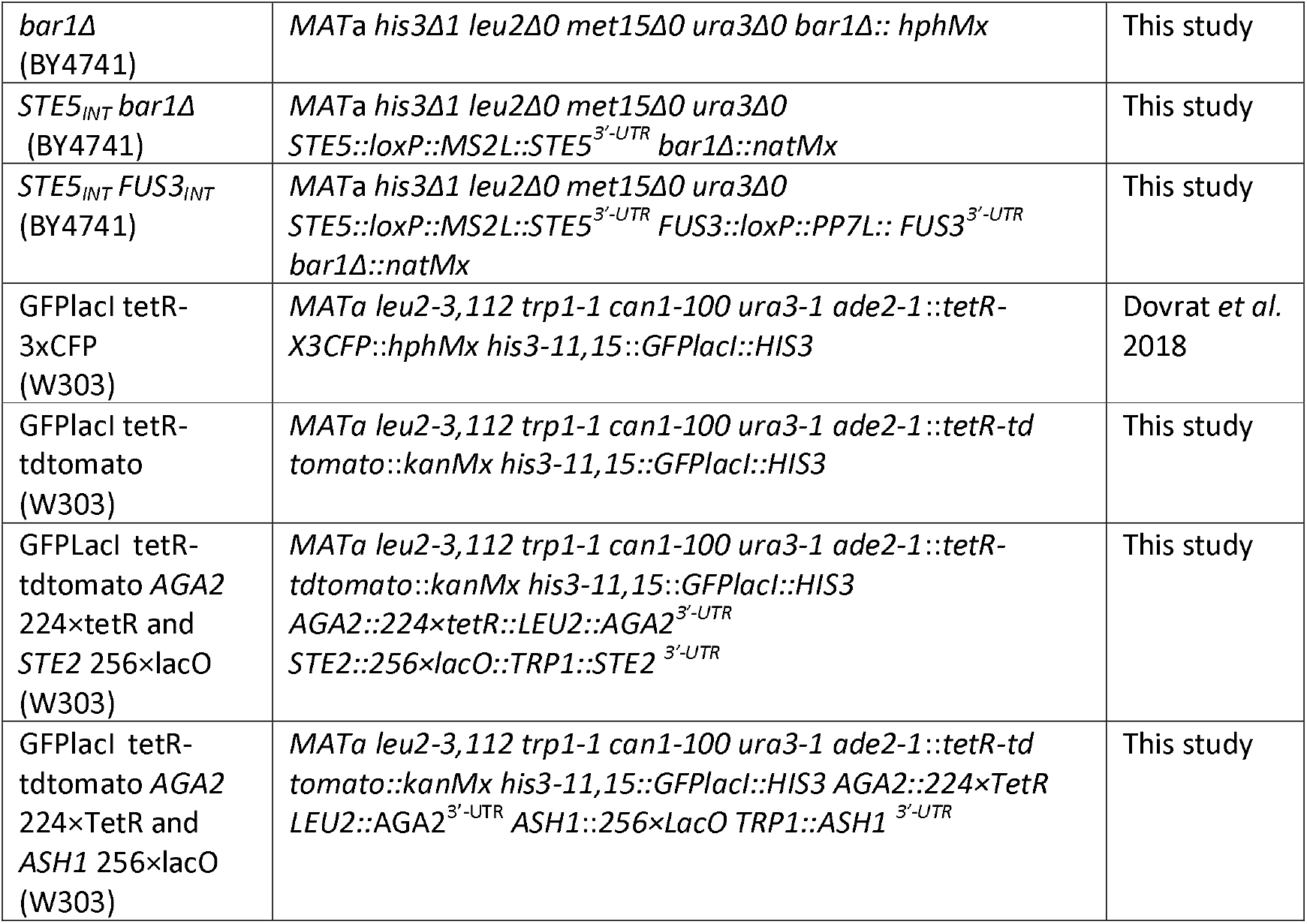
Yeast strains used in this study.

**Table S3.** Plasmids used in this study.

| Plasmid name | Gene expressed | Vector | Copy number | Selection marker | Source |
| --- | --- | --- | --- | --- | --- |
| pMS2-CP-GFPx3 | *MET25p-MS2-CP-GFP(x3)* | pCP-GFP (pUG23 base) | *CEN* | *HIS3* | J. Gerst |
| pMS2-CP-GFPx3 | *MET25p-MS2-CP-GFP(x3)* | pCP-GFP (pUG23 base) | *CEN* | *URA3* | J. Gerst |
| pAD54-Hhf1 | *HA-HHF1* | pAD54 | 2μ | *LEU2* | This Study |
| pAD4Δ-Hhf1-RFP | *HHF1-RFP* | pAD4Δ | 2μ | *LEU2* | This Study |
| prs426-Cas9-sgDNA-natMx | Cas9 sgRNA against *natMx* marker | pRS426 | 2μ | *URA3* | This Study |
| prs426-Cas9-sgDNA-HO | Cas9 sgRNA against *HO* gene | pRS426 | 2μ | *URA3* | This Study |
| pmCherry-Scs2 | *mCherry-SCS2* | pAD4Δ | 2μ | *LEU2* | J. Gerst |
| pLOXHIS5MS2L | *MS2L, Sphis5+* | pUG27 | *CEN* | *Sphis5+* | J. Gerst |
| pSH47 | *GALp-CRE* | - | *CEN* | *URA3* | Euroscarf |
| pAD54-Hhf1<sup>K-R</sup> | *HA-HHF1<sup>K-R</sup>* | pAD54 | 2μ | *LEU2* | This Study |
| pAD54-Hhf1<sup>K-Q</sup> | *HA-HHF1<sup>K-Q</sup>* | pAD54 | 2μ | *LEU2* | This Study |
| pFA6-natNT2 | - | - | - | *natMx (NAT)* | Euroscarf |
| pFA6-hphMx | - | - | - | *hphMx (hygromycin B)* | Euroscarf |
| pRS316-AGA2 | *AGA2* | pRS316 | *CEN* | *URA3* | This Study |
| pRS316-AGA2 27mut | *AGA2 27 RBM mut* | pRS316 | *CEN* | *URA3* | This Study |
| pRS316-AGA2 47mut | *AGA2 47 RBM mut* | pRS316 | *CEN* | *URA3* | This Study |
| pSR10 | *lacOx256 array* | - | - | *TRP1* | S. Gasser |
| pDtet | *tetOx224* | - | - | *LEU2* | A. Aharoni |
| pDtetR-tdTomato | *tetR-tdTomato* | - | - | *kanMx (Kanamycin)* | A.Aharoni |

